# Marigold: A machine learning-based web app for zebrafish pose tracking

**DOI:** 10.1101/2024.05.31.596910

**Authors:** Gregory Teicher, R. Madison Riffe, Wayne Barnaby, Gabrielle Martin, Benjamin E. Clayton, Josef G. Trapani, Gerald B. Downes

## Abstract

High-throughput behavioral analysis is important for drug discovery, toxicological studies, and the modeling of neurological disorders such as autism and epilepsy. Zebrafish embryos and larvae are ideal for such applications because they are spawned in large clutches, develop rapidly, feature a relatively simple nervous system, and have orthologs to many human disease genes. However, existing software for video-based behavioral analysis can be incompatible with recordings that contain dynamic backgrounds or foreign objects, lack support for multiwell formats, require expensive hardware, and/or demand considerable programming expertise. Here, we introduce Marigold, a free and open source web app for high-throughput behavioral analysis of embryonic and larval zebrafish. Marigold features an intuitive graphical user interface (GUI), tracks up to 10 user-defined keypoints, supports both single- and multiwell formats, and exports a range of kinematic parameters in addition to publication-quality data visualizations. By leveraging a highly efficient, custom-designed neural network architecture, Marigold achieves reasonable training and inference speeds even on modestly powered computers lacking a discrete graphics processing unit (GPU). Notably, as a web app, Marigold does not require any installation and runs within popular web browsers on ChromeOS, Linux, macOS, and Windows. To demonstrate Marigold’s utility, we conducted two sets of biological experiments. First, we examined novel aspects of the touch-evoked escape response in *techno trousers (tnt)* mutant embryos, which contain a previously described loss-of-function mutation in the gene encoding Eaat2b, a glial glutamate transporter. We identified differences and interactions between touch location (head vs. tail) and genotype. Second, we investigated the effects of feeding on larval visuomotor behavior at 5 and 7 days post-fertilization (dpf). We found differences in the number and vigor of swimming bouts between fed and unfed fish at both time points, as well as interactions between developmental stage and feeding regimen. In both biological experiments presented here, the use of Marigold facilitated novel behavioral findings. Marigold’s ease of use, robust pose tracking, amenability to diverse experimental paradigms, and flexibility regarding hardware requirements make it a powerful tool for analyzing zebrafish behavior, especially in low-resource settings such as course-based undergraduate research experiences (CUREs). Marigold is available at: https://downeslab.github.io/marigold/.

## Introduction

The kinematic and ethological analysis of animal movements is an essential task for many researchers in neuroscience, psychology, and related fields. For investigators working with zebrafish (*Danio rerio*), behavioral assays are key methods for characterizing the toxicological effects of environmental pollutants, identifying therapeutic agents using small molecule screens, and modeling neurological disorders such as autism and epilepsy (Baraban et al., 2013; d’Amora & Giordani, 2018; MacRae & Peterson, 2023; Sakai et al., 2018). Zebrafish embryos are spawned in large clutches and develop rapidly, making high-throughput screening in multiwell plates an attractive experimental paradigm. Additionally, zebrafish have a well characterized behavioral repertoire, a relatively simple nervous system containing fewer neurons compared to many other vertebrates, and orthologs to numerous human disease genes (Berg et al., 2018; Holtzman et al., 2016; Howe et al., 2013; Kalueff et al., 2013). Thus, zebrafish are an excellent system for studying vertebrate behavior.

Despite the rising popularity of behavioral screening in zebrafish, currently available software for analysis has a variety of limitations. Behavioral analysis software has traditionally utilized classical image processing techniques such as pixel intensity thresholding, background subtraction, skeletonization, and center of mass calculations to isolate animals from the visual background and track their movements (Burgess & Granato, 2007a; Clift et al., 2014; Joo et al., 2020; Marques et al., 2018; Mirat et al., 2013). These techniques are often sufficient, but can lack robustness when challenged with dynamic backgrounds, varying lighting conditions, or earlier developmental stages when embryos present with less contrast against the background. Many solutions provide relatively limited kinematic information by only tracking a single point representing a larva’s overall position, while those programs capable of tracking multiple points often fail when foreign objects enter the field of view. Approaches based on classical image processing techniques are particularly problematic for touch-evoked response assays that require a probe to enter the field of view, especially when the tactile stimulus is applied to the tail rather than the more easily detectable head. Additionally, many programs are closed source and/or tied to proprietary hardware such as specialized imaging cabinets, thus restricting access and customizability.

Several recently introduced programs have extended machine learning-based advances in human pose tracking to other species (Goodwin et al., 2024; Gore et al., 2023; Graving et al., 2019; Mathis et al., 2018; Pereira et al., 2019; Ravan et al., 2023; Romero-Ferrero et al., 2019). Such programs can achieve high accuracy, but they commonly require a high-end graphics processing unit (GPU) to run the large deep learning models efficiently. Additionally, many of these programs require complex workflows combining GUI-, command line-, and cloud-based elements, making such software difficult to use for those who lack programming, machine learning, or other technical expertise. These factors also limit the use of such programs in resource-constrained settings such as course-based undergraduate research experiences (CUREs), where such software might otherwise be useful. Lastly, due to the general-purpose nature of existing deep learning-based pose tracking programs, they often lack support for popular zebrafish experimental paradigms, such as imaging in multiwell plates, which can limit their utility for zebrafish researchers.

Here, we present Marigold, a free and open source machine learning-based web app for automated pose tracking of embryonic and larval zebrafish that unites many of the strengths of existing programs, while also addressing several common limitations. Marigold reliably tracks up to 10 user-defined body keypoints, supports both single- and multi-well configurations, and features a user-friendly interface (Figure 1). Marigold facilitates a streamlined workflow consisting of three stages: (1) generating a dataset by extracting a modest number of frames from representative behavioral recordings and labeling the frames with animal pose coordinates; (2) training a neural network using the generated dataset; and (3) analyzing new behavioral recordings using the trained neural network and exporting the resulting raw kinematic parameters and visualizations. Despite using convolutional neural networks, which are notoriously computationally demanding, Marigold is able to train models and analyze behavioral recordings at reasonable speeds on current desktop or laptop computers without requiring a dedicated GPU. This speed is achieved by using a highly efficient, custom neural network architecture incorporating a series of macro-architectural and micro-architectural optimizations. For each animal analyzed, Marigold generates publication-ready trajectory plots and a CSV file of keypoint coordinates, speeds, and angles between keypoints, which facilitates downstream analysis pipelines. Notably, as a web app, Marigold does not require any installation and is highly cross-platform, running within popular web browsers on ChromeOS, Linux, macOS, and Windows across a wide range of devices.

**Figure 1.**
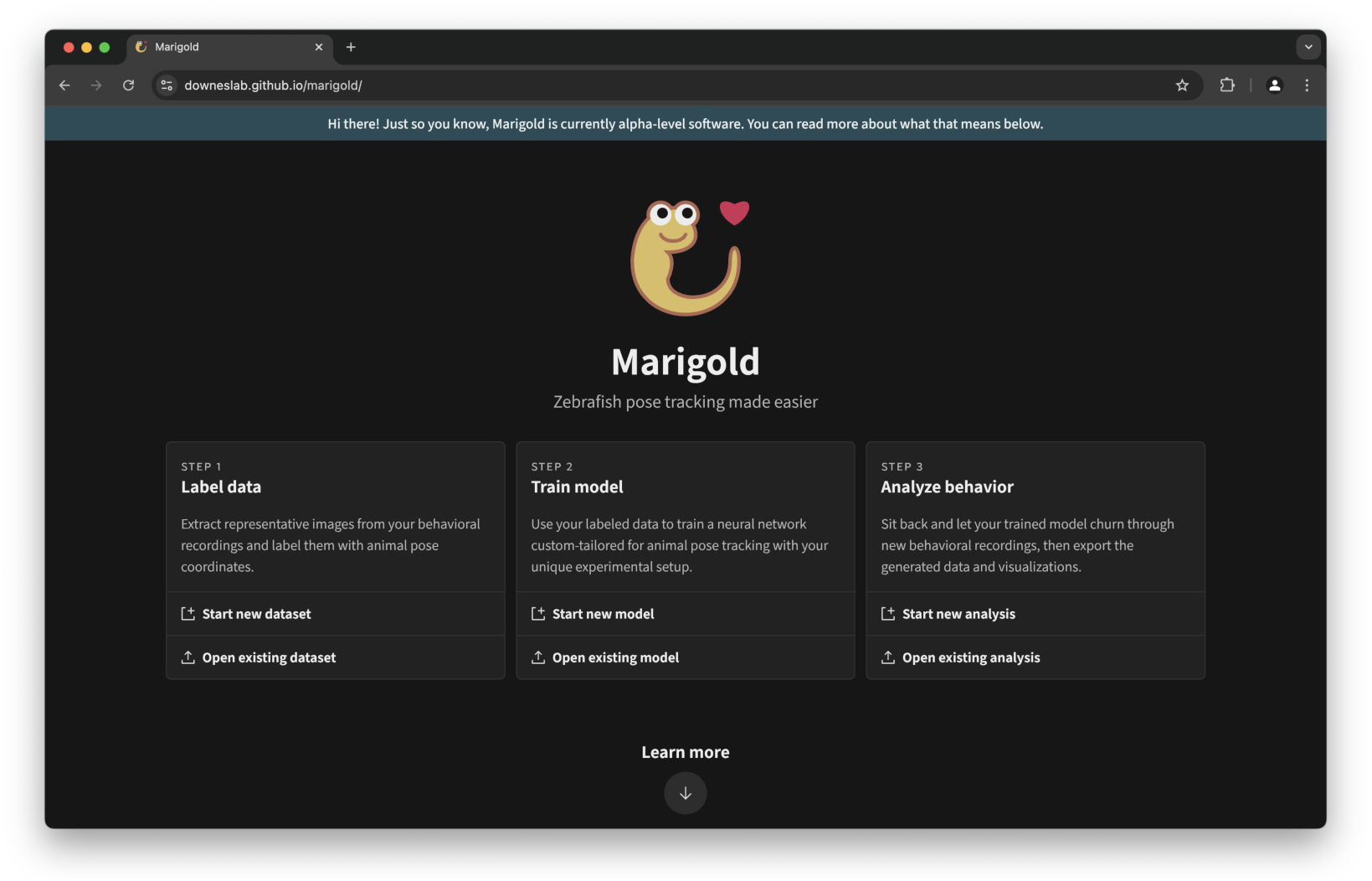
Marigold features an intuitive web-based graphical user interface (GUI). Marigold’s welcome page (as viewed in Google Chrome on macOS) orienting the user to the workflow of labeling a dataset, training a model, and analyzing behavior.

To showcase its robustness and versatility, we used Marigold to obtain biological insights from two sets of zebrafish experiments. First, we examined novel aspects of the embryonic touch-evoked response phenotype in the previously described *techno trousers (tnt)* mutant (Granato et al., 1996; McKeown et al., 2012), which contains a loss-of-function mutation in the gene encoding Eaat2b, a glial glutamate transporter. By applying the stimulus to either the head or tail, we identify differences and interactions between genotype and touch location, potentially providing new clues into how neural circuits may be disrupted in this mutant. Second, we investigated the effects of developmental stage and feeding on larval visuomotor response behavior. We identify differences and interactions between feeding regimen and developmental stage, highlighting the importance of considering both factors when designing and reporting the results of larval visuomotor response assays. In both biological experiments, the use of Marigold led to novel behavioral findings, demonstrating some of the types of biological questions that can be addressed using the detailed kinematic analysis provided by Marigold.

## Methods

### Zebrafish husbandry

All animals used in this study were obtained from our lab’s fish facility at University of Massachusetts Amherst. Adult zebrafish were bred to obtain embryos and larvae for experimentation. Embryos and larvae younger than 5 dpf were maintained in the dark at 28.5 °C. Older fish were maintained on a 14 h light, 10 h dark cycle at approximately 28.5 °C. All wild type fish were from either Tübingen or Tüpfel long-fin (TL) strains as indicated. *techno trousers* (*slc1a2b^tk57^*) fish (Granato et al., 1996; McKeown et al., 2012) were maintained on a TL background. All animal procedures were approved by the University of Massachusetts Amherst Institutional Animal Care and Use Committee (IACUC) under Animal Welfare Assurance number A3551-01. No animal procedures were performed at Amherst College.

### Dataset collection and labeling

Two novel datasets were developed: one for touch-evoked responses and one for visuomotor responses. For both datasets, embryos were obtained from mass matings of Tübingen or TL fish. Behavior was monitored using an IDT X-Stream 1440p PCIe 2.0 high speed camera (Integrated Design Tools, Inc., Pasadena, CA, USA). Larvae were imaged from above and illuminated using a white light or near-infrared source from below. Recordings were made under varying conditions of lighting intensity, resolution, magnification, and focus. From the generated movies, frames were manually selected for inclusion in the dataset with the goal of maximizing the representation of diverse poses and experimental conditions. Each frame was then manually labeled and the labeled coordinates were used to generate target heatmaps (i.e., two-dimensional Gaussians) that served as the “ground truth” for our neural networks during training.

For the touch-evoked response dataset, movies were recorded of embryos and larvae between the developmental stages of 2–4 dpf in wells in the lids of 24-well plates. Manual labels targeted 7 keypoints evenly spaced across the rostral caudal axis. For the visuomotor response dataset, movies were recorded of larvae between the developmental stages of 5–7 dpf in 24-well plates. Manual labels targeted a single keypoint positioned between the two eyes.

### Software development

Marigold was written in HTML, CSS, JavaScript, and C++ (with the latter used to generate WebAssembly) and is licensed under the GNU General Public License version 3 (GPLv3). No external libraries or components were used or incorporated other than the Adobe Source Sans 3 font (used under the SIL Open Font License Version 1.1) (Adobe, n.d.), the plasma and viridis colormaps (used under the CC0 “No Rights Reserved” license) (Smith et al., n.d.), and the SplitMix64 (Steele et al., 2014) and xoshiro256** (Blackman & Vigna, 2021) pseudorandom number generators (both used under the CCO “No Rights Reserved” license) (Blackman & Vigna, n.d.). Building for deployment was automated using GitHub Actions, with hosting provided by GitHub Pages.

### Measurement and calculation of performance metrics

Performance measurements were obtained using a Lenovo Thinkpad P14s Gen 4 laptop equipped with an AMD Ryzen 7 PRO 7840U central processing unit (CPU) and running Fedora Workstation 41 with Linux kernel version 6.11.7. PyTorch-based performance measurements were obtained using Python 3.13.0, NumPy 2.1.3 (Harris et al., 2020), pandas 2.2.3 (McKinney, 2010), SciPy 1.14.1 (Virtanen et al., 2020), PyTorch 2.5.1 (Ansel et al., 2024), and torchvision 0.20.1 (TorchVision maintainers and contributors, 2016). WebAssembly-based performance measurements were obtained using the LLVM (version 19.1.0) Clang compiler and wasm-ld linker, with the generated WebAssembly running in Google Chrome 131. All performance measurements reflect single-threaded CPU performance.

Training speeds reflect the time required to complete a forward and backward pass through the neural network in training mode at the specified batch size. Inference speeds reflect the time required to complete a forward pass through the neural network in evaluation mode at the specified batch size. Notably, these measurements do not include the time required to perform data augmentation or gather data into batches.

Theoretical minimum memory footprints correspond to a hypothetical implementation in which all required memory buffers must be allocated before training begins and cannot be freed or repurposed until training has ended, individual computations are executed with the minimum possible number and size of memory buffers (e.g., by using implicit rather than explicit padding for depthwise convolutions), and gradient accumulation is used to the fullest extent possible wherever it can be used without impacting the results (i.e., in the absence of Batch Normalization).

### Data augmentation

Datasets were randomly divided into training (75%) and validation (25%) splits. On-the-fly data augmentation (Shorten & Khoshgoftaar, 2019) was applied to images in the training split by randomly adjusting gamma and brightness, randomly resizing, randomly flipping vertically and horizontally, randomly rotating, randomly padding and cropping, and (after standardization) randomly adding a small amount of Gaussian noise. Additionally, a small amount of “wiggle” was randomly applied to keypoint coordinates during generation of the two-dimensional gaussian labels. Data augmentation was not applied to images or labels in the validation split. All images were standardized using the mean and standard deviation calculated over the training split.

### Neural network design and training

Neural networks were trained at an input resolution of 512×512 (touch-evoked response dataset) or 256×256 (visuomotor response dataset). All MobileNetV3 blocks and variations used an expansion ratio of 2, with the number of channels as input to the block set to 32 (touch-evoked response dataset) or 16 (visuomotor response dataset) when operating at the smallest spatial resolution, except where otherwise noted. For the hierarchical macro-architecture, intro blocks consisted of a 4×4 convolution with 2×2 stride followed by a normalization layer. Outro blocks consisted of a 1×1 convolution layer with a channel expansion ratio of 2, a hard swish layer, a spatial dropout layer with drop probability = 0.05, and a 1×1 convolution layer reducing the channels to the number of keypoints. Downsampling and upsampling blocks consisted of a 2×2 convolution with 2×2 stride or a 2×2 transposed convolution with 2×2 stride stride followed by a normalization layer. For the isotropic macro-architecture, intro blocks consisted of a 8×8 pixel unshuffle layer, a 1×1 convolution layer, and a normalization layer, except where otherwise noted. Outro blocks consisted of a 1×1 convolution layer with a channel expansion ratio of 2, a hard swish layer, a spatial dropout layer with drop probability = 0.05, a 1×1 convolution layer reducing the channels to 16× the number of keypoints, and a 4×4 pixel shuffle layer transforming the number of channels to the number of keypoints while increasing the spatial resolution, except where otherwise noted.

Training was performed using mean squared error (MSE) loss, a batch size of 16, and the AdamW optimizer (Loshchilov & Hutter, 2019) with learning rate = 5.0 × 10^−4^ (touch-evoked response dataset) or 1.0 × 10^−4^ (visuomotor response dataset), β_1_ = 0.9, β_2_ = 0.95, and ε = 1.0 × 10^−6^. Untuned learning rate warmup was used, as suggested by Ma and Yarats, 2021. Weight decay = 1.0 × 10^−5^ was applied to the weights of all convolution layers. Convolution weights were initialized using Xavier initialization (Glorot & Bengio, 2010), with the exception of the final convolution layer which was initialized to zeros. When a convolution layer was followed by a nonlinearity or by a normalization and then nonlinearity, the initialization gain was set to the square root of two, with the gain otherwise set to one. Additionally, the initialization gain for the last convolution layer in each residual block was scaled by the reciprocal of the square root of the number of residual blocks, as suggested by Marion et al., 2024. All convolution biases were initialized to zeros. For all normalization layers, ε was set to 1.0 × 10^−3^, which matches the value used for Batch Normalization in MobileNetV3 (A. Howard et al., 2019) and was found to have a stabilizing effect on training compared to the PyTorch default value of 1.0 × 10^−5^. The PyTorch default value of momentum = 0.9 was used for Batch Normalization layers. Group Normalization was configured to use 4 channels per group. Layer Normalization and Instance Normalization were implemented in PyTorch as Group Normalization with one group or as many groups as channels, respectively.

### Touch-evoked response assay

Embryos were obtained from mass matings of heterozygous *techno trousers* (*slc1a2b^+/tk57^*) fish. Behavior was monitored using an IDT X-Stream 1440p PCIe 2.0 high speed camera (Integrated Design Tools, Inc., Pasadena, CA, USA) in a temperature controlled setting (25 °C). Larvae were transferred in E3 solution to a well in the inverted lid of a CytoOne 24-Well Tissue Culture Plate (USA Scientific, Inc., Ocala, FL, USA) and allowed to acclimate briefly before touching on either the side of the head or tip of the tail as indicated with a 3.22/0.16 g of force von Frey filament held by a surgical blade holder. Larvae were imaged from above and illuminated using a white light source from below. Responses were recorded at a resolution of 1024×1024 pixels with a framerate of 1000 Hz and exposure time of 0.8 ms. Recordings were manually trimmed to remove extraneous frames before and after the response, as determined by the first and last visible movements initiated by the fish, respectively.

### *slc1a2b* genotyping

Following behavioral analysis, embryos were euthanized using an overdose of MS-222 (pH = 7.0) (Sigma-Aldrich, St. Louis, MO, USA). DNA was extracted from the euthanized embryos using Extract-N-Amp Tissue PCR Kit (Sigma-Aldrich, St. Louis, MO, USA) according to the manufacturer’s protocol. A 192 bp region of exon 2 of *slc1a2b* flanking the site encoding the A393V mutation was amplified by PCR using the forward primer 5’-TGCTGGAACTCTGCCCGTGA-3’ and the reverse primer 5’-ACGGTGACGATCTGTCCAGG-3’.

PCR was performed using AmpliTaq Gold DNA Polymerase (Applied Biosystems/Thermo Fisher Scientific, Waltham, MA, USA) according to the manufacturer’s protocol and using an annealing temperature of 59 °C. The PCR product was digested overnight using Fnu4HI (New England Biolabs, Ipswich, MA, USA) according to the manufacturer’s protocol. Digestion yielded a single fragment of 192 bp for homozygous mutant fish (*slc1a2b^tk57/tk57^*), two fragments of 133 and 59 bp for homozygous wild type fish (*slc1a2b^+/+^*), and three fragments of 192, 133, and 59 bp for heterozygous fish (*slc1a2b^+/tk57^*). The digested PCR products were resolved for genotypic determination by agarose gel electrophoresis.

### Visuomotor response assay

Embryos were obtained from mass matings of Tübingen strain zebrafish and maintained in E3 media for the duration of experiments. Larvae were kept in petri dishes until 5 dpf, at which point they were transferred to 2.5 L fish tanks at a density of approximately 45 fish/tank and maintained on a 14 h light, 10 h dark light cycle. The initial volume of E3 media in the tanks was 250 mL, with an additional 250 mL added daily. Fish in fed conditions were given approximately 50 mg of Gemma Micro 75 (Skretting, Stavanger, Norway) once daily beginning at 5 dpf and continuing through 7 dpf. Feeding occurred at approximately 12:00 PM each day, with behavioral recordings occurring between 4-6 h after feeding. Behavior was monitored using an IDT X-Stream 1440p PCIe 2.0 high speed camera (Integrated Design Tools, Inc., Pasadena, CA, USA) in a temperature controlled setting (25 °C). Larvae were transferred to the wells of a CytoOne 24-Well Tissue Culture Plate (USA Scientific, Inc., Ocala, FL, USA) for recording. Larvae were imaged from above and illuminated from below using a custom-built light source which was toggled between white and near-infrared illumination modes. Fish were shielded from ambient light by an enclosing cabinet. Responses were recorded at a resolution of 1440×1440 pixels with a framerate of 100 Hz and exposure time of 0.8 ms.

### Statistical analysis and data visualization

Statistical significance was determined by two-way analysis of variance (ANOVA) with Tukey post hoc test using the R language (version 4.4.2) (R Core Team, 2024). Data visualizations were generated using the ggplot2 library (version 3.5.1) for R (Wickham, 2016).

## Results

### Neural network design

Arguably, modern machine learning is dominated by the massively parallel training of extremely overparameterized deep neural networks on colossal datasets using high-end GPU clusters (Greener et al., 2022; Togelius & Yannakakis, 2024). This paradigm is at odds with common use case scenarios in resource-constrained settings such as course-based undergraduate research experiences (CUREs) and academic life science research labs, in which hardware often lags behind the cutting edge and domain-specific datasets frequently must be collected and labeled *de novo*.

To reduce the processing and memory demands of deep convolutional neural networks sufficiently to make in-browser, on-device zebrafish pose tracking feasible with minimal hardware requirements and limited data availability, we collected and labeled two proof-of-principle datasets and used them to develop and evaluate a series of neural network architectural variants. Our datasets were designed to reflect behavioral assays commonly used within our lab and were composed of either labeled frames from touch-evoked response movies, in which case labels consisted of 7 keypoints evenly spaced across the rostral–caudal axis, or labeled frames from visuomotor response movies, in which case labels consisted of a single keypoint located between the two eyes (Figure 2). Our neural networks were prototyped using PyTorch, a highly optimized machine learning library for Python, and evaluated based on training dynamics, training speed, inference speed, parameter count, and theoretical minimum memory footprint. We chose to evaluate training and inference speeds primarily based on single-threaded CPU performance as this represents a “lowest common denominator” target across the wide variety of platforms, devices, and research settings we aimed to support.

**Figure 2.**
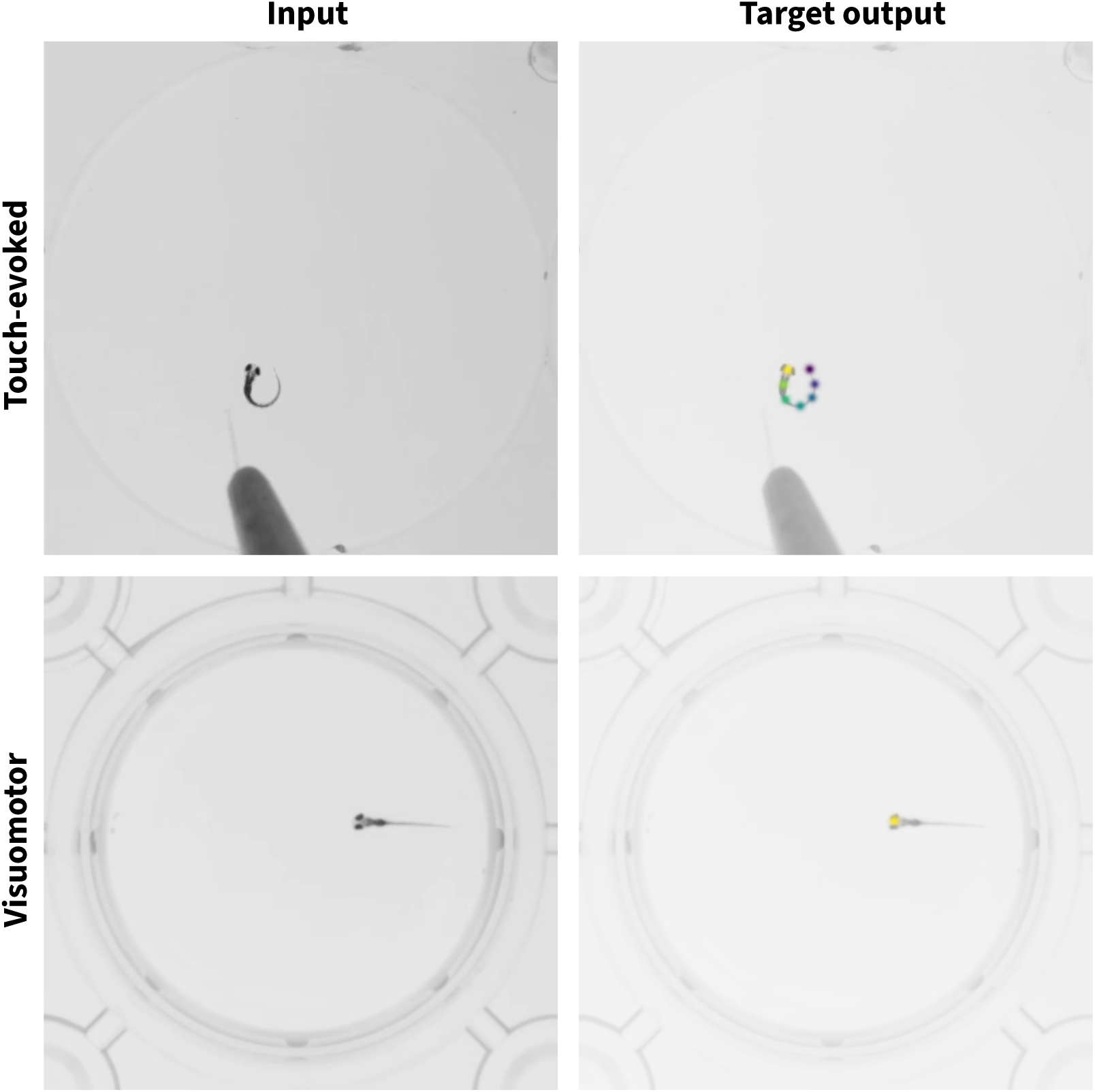
Examples of input–output pairs from the touch-evoked response and visuomotor response datasets. Representative images from each dataset visualized with and without overlaid pseudocolored gaussians corresponding to labeled keypoints.

As an efficient and well-established baseline building block for our neural network architectural design experiments, we adopted the MobileNetV3 inverted bottleneck residual block (A. Howard et al., 2019). As the most recent member of the MobileNet family of blocks designed for fast CPU inference on mobile devices (A. Howard et al., 2019; A. G. Howard et al., 2017; Sandler, Howard, et al., 2019), the MobileNetV3 design improved on that of MobileNetV2 by optionally incorporating larger depthwise convolution kernel sizes, the Hard Swish activation function, and the Squeeze and Excitation module.

Having selected the MobileNetV3 inverted bottleneck residual block as our baseline building block, we proceeded to incorporate it into two potential macro-architectural designs (Figure 3A). In the first approach, we adopted a hierarchical architecture inspired by networks such as the Hourglass and Simple Baseline architectures for human pose estimation and the U-Net architecture for semantic segmentation of medical images (Newell et al., 2016; Ronneberger et al., 2015; Xiao et al., 2018). In this architecture, as information flows through the successive layers of the network, it is first downsampled, then upsampled. The downsampling path minimizes the computational cost of the expensive convolutional layers and aggregates spatial context, while the upsampling path restores the resulting representation to the larger spatial resolution required to produce accurate probability heatmaps for each body keypoint of interest. In the second approach, we adopted an isotropic architecture (Sandler, Baccash, et al., 2019). In contrast to hierarchical architectures which make use of multiple internal resolutions, isotropic architectures utilize a constant internal resolution, essentially performing all processing on non-overlapping image “patches”. Isotropic architectures also remain relatively unexplored, particularly for computer vision tasks other than image classification. Interestingly, an isotropic architecture was recently used for keypoint-based fish morphometric analysis (Saleh et al., 2023), suggesting its potential utility for animal pose estimation. We tailor our architectures to our two datasets as described in Table 1.

**Figure 3.**
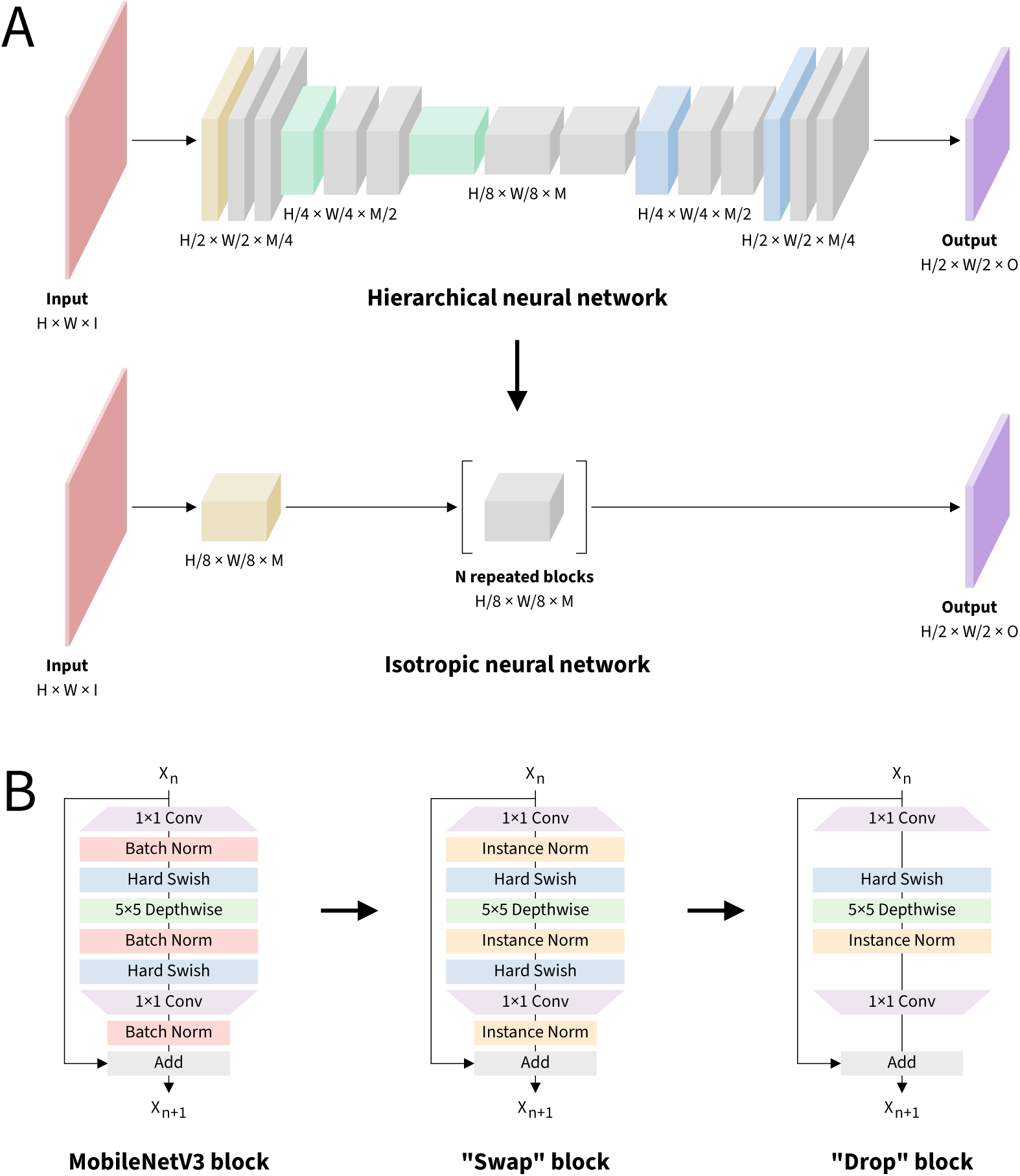
Overview of macro-architectural and micro-architectural design decisions leading to a simple, efficient, and easily customizable neural network architecture. (**A**) Schematic of the macro-architectural design process illustrating the implementations of hierarchical and isotropic neural networks. Red indicates input images, yellow indicates intro blocks, green indicates downsampling blocks, blue indicates upsampling blocks, purple indicates outro blocks, and gray indicates residual blocks. H: input height; W: input width; I: input channels; M: middle channels; O: output channels; N: residual blocks. (**B**) Schematic of the microarchitectural design process illustrating the implementations of the MobileNetV3 inverted bottleneck residual block and subsequent modifications. Small arrows indicate the flow of information through the macro- and micro-architectures, while large arrows indicate the flow of the design process.

**Table 1.** Neural network configurations. Designations in parentheses correspond to those in Figure 3A.

| Dataset | Touch-evoked | Visuomotor |
| --- | --- | --- |
| Input resolution ($H \times W$) | $512 \times 512$ | $256 \times 256$ |
| Output resolution ($H/2 \times W/2$) | $256 \times 256$ | $128 \times 128$ |
| Input channels (I) | 1 | 1 |
| Middle channels (M) | 32 | 16 |
| Output channels (O) | 7 | 1 |
| Residual blocks (N) | 10 | 10 |

Despite its simplicity, the isotropic architecture exhibited compelling advantages over its hierarchical counterpart (Figure 4). Across both datasets, the isotropic architecture reached lower loss values in less time (Figure 4A–B) and achieved severalfold faster training and inference throughput (Figure 4C–D). Although the isotropic architecture contains a greater number of trainable parameters, its theoretical minimum memory footprint is considerably smaller (Figure 4E–F). To help tease apart any intrinsic benefit of patch-based processing from that of operating at a specific internal resolution, we also explored using patches larger and smaller than the 8×8 patches corresponding to the smallest internal resolution of our hierarchical architecture, adjusting the number of channels in inverse proportion to the number of spatial pixels to control for the overall processing and memory demands. Smaller patch sizes led to considerably slower training convergence while larger patch sizes led to similar training convergence but with lower inference speed; thus, 8×8 patches appeared to provide the most optimal balance across our metrics of interest (Supplementary Figure 1). Altogether, moving from a hierarchical architecture to an isotropic architecture appears to be a net positive design decision.

**Figure 4.**
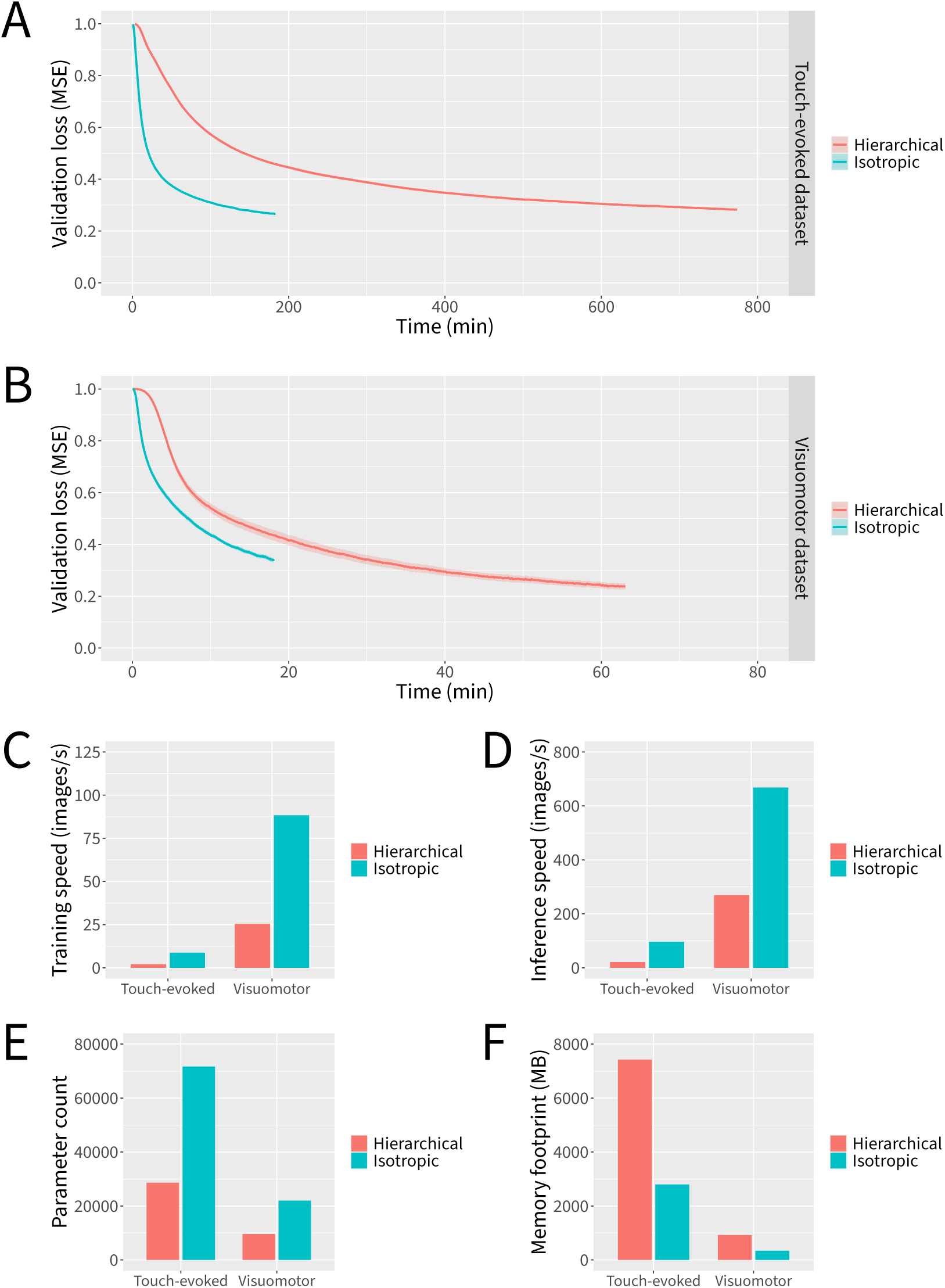
Isotropic neural network macro-architectures outperform more traditional hierarchical neural network macro-architectures by multiple metrics. Neural networks were trained for 6000 parameter updates for each dataset and macro-architecture. (**A**) Loss curves for the touch-evoked response dataset. (**B**) Loss curves for the visuomotor response dataset. (**C**) Training speeds. (**D**) Inference speeds. (**E**) Parameter counts. (**F**) Theoretical minimum memory footprints. Data represent the mean or the mean plus and minus the standard error of the mean of 10 independent experiments in which datasets were partitioned and neural network weights were initialized using different random seeds.

With our macro-architectural design strategy in place, we turned our attention to micro-architectural improvements to the MobileNetV3 block (Figure 3B; Figure 5). As our first micro-architectural design step, we explored alternatives to the Batch Normalization layers used in the MobileNetV3 block. Batch Normalization is widely adopted in neural network architectures for computer vision due to its ability to improve training dynamics and help prevent overfitting (Ioffe & Szegedy, 2015). During training, the computational overhead introduced by Batch Normalization leads to slower per-epoch training throughput, but this is generally outweighed by the resulting improvement in per-epoch training convergence. At inference time, the computational overhead can be avoided by “folding” Batch Normalization layers into adjacent convolutional layers. Thus, Batch Normalization generally leads to more efficient training without negatively impacting inference speed. However, the dependence of Batch Normalization on the batch size used during training leads on the one hand to markedly poorer training convergence at smaller batch sizes and, on the other hand, to a larger training memory footprint at large batch sizes, with the potential to exceed the limited amount of memory that web apps are permitted to allocate. By contrast, proposed alternatives to Batch Normalization, such as Layer Normalization (Ba et al., 2016), Group Normalization (Wu & He, 2018), and Instance Normalization (Ulyanov et al., 2017), operate independently of batch size, enabling the use of gradient accumulation to maintain a constant training memory footprint independent of batch size. However, these alternatives cannot be “folded” into preceding convolutional layers, introducing computational overhead at inference time. In our experiments, Layer Normalization, Group Normalization, and Instance Normalization all led to competitive training convergence compared to Batch Normalization at a modest cost to inference speed (Supplementary Figure 2). However, “swapping” from Batch Normalization to Instance Normalization resulted in the greatest improvement in training dynamics across both datasets and thus was incorporated into our final architectural design (Figure 3B; Figure 5).

**Figure 5.**
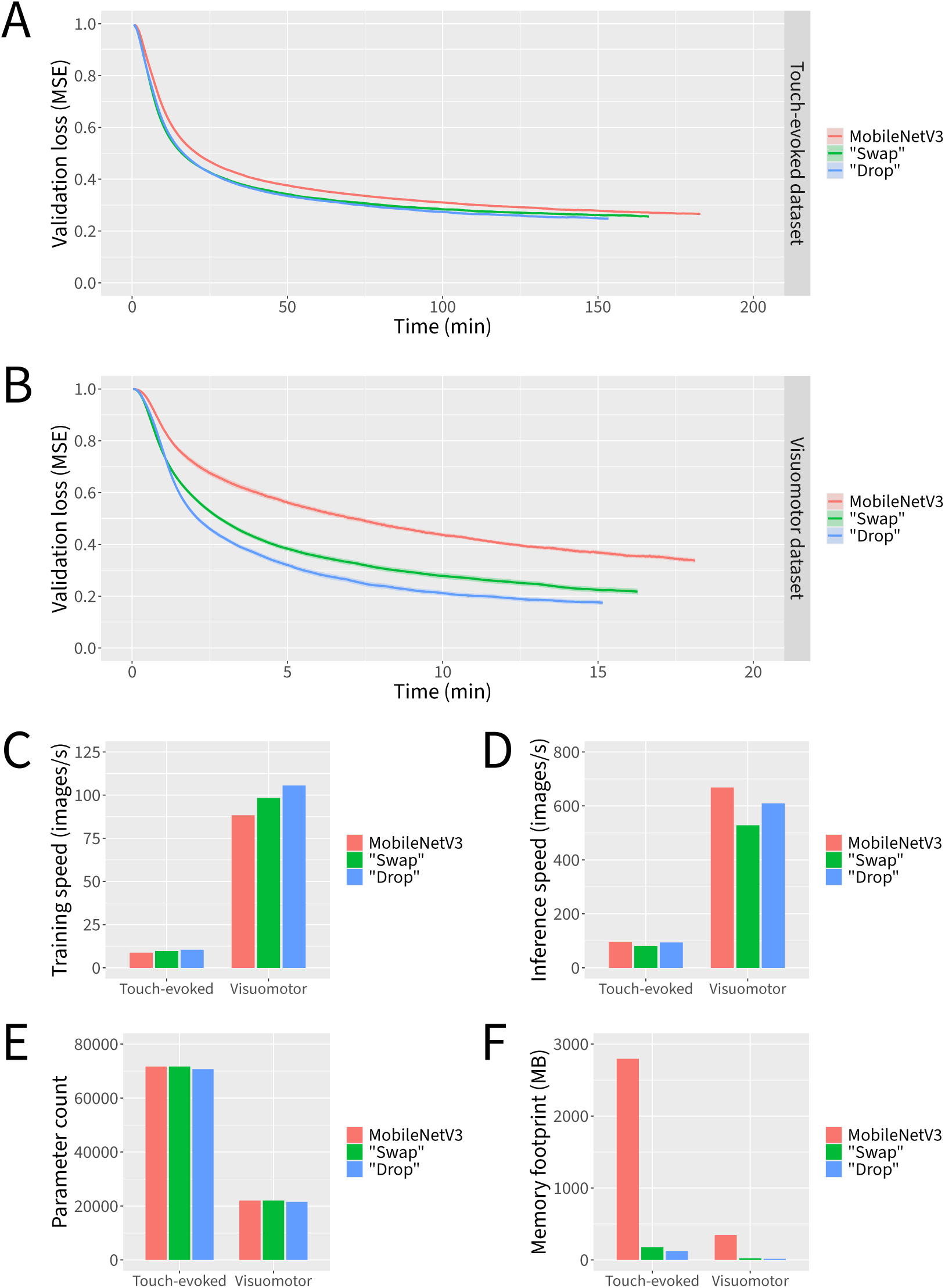
Micro-architectural improvements to the MobileNetV3 block improve training dynamics and reduce memory requirements at minimal cost to inference speed. Neural networks were trained for 6000 parameter updates for each dataset and micro-architecture. (**A**) Loss curves for the touch-evoked response dataset. (**B**) Loss curves for the visuomotor response dataset. (**C**) Training speeds. (**D**) Inference speeds. (**E**) Parameter counts. (**F**) Theoretical minimum memory footprints. Data represent the mean or the mean plus and minus the standard error of the mean of 10 independent experiments in which datasets were partitioned and neural network weights were initialized using different random seeds.

Next, we asked whether it was possible to recover some of the inference speed that was lost as a result of the switch from Batch Normalization to Instance Normalization without negatively impacting the training dynamics (Figure 3B). A logical way to accomplish this would be to “drop” some of the components of the MobileNetV3 block. We thus performed a series of ablation studies to determine which components were most dispensable. We were surprised to find that “dropping” two of the three Instance Normalization layers and one of the two Hard Swish layers actually led to more favorable training dynamics than any other design iteration, while also recovering much of the inference speed (Figure 5). We note that the choices of which Instance Normalization layer and Hard Swish layer to keep were crucial in this design step, as other combinations did not perform as well (Supplementary Figure 3).

Altogether, our series of macro-architectural and micro-architectural design steps resulted in markedly faster training and inference speeds, with substantially reduced theoretical memory footprints, compared to our already reasonably efficient baseline. Given that similar results were obtained across both datasets, our final architecture appears to be flexible enough to accommodate a range of experimental paradigms. All the more so considering that the isotropic nature of our architecture largely decouples design parameters such as the input resolution, middle resolution, output resolution, number of middle channels, number of residual blocks, and residual block channel expansion ratio, thus supporting a large degree of potential customization.

### Training data requirements

Given that manually labeling data is often the most labor-intensive aspect of any supervised training workflow, we next asked to what extent similar results could be obtained with less training data available. We investigated this by incrementally ablating images from the training split of each of our two datasets while controlling for the total number of parameter updates (Figure 6). To preserve our ability to fairly evaluate each model’s ability to generalize to previously unseen data, we held the number of images in the validation split constant.

**Figure 6.**
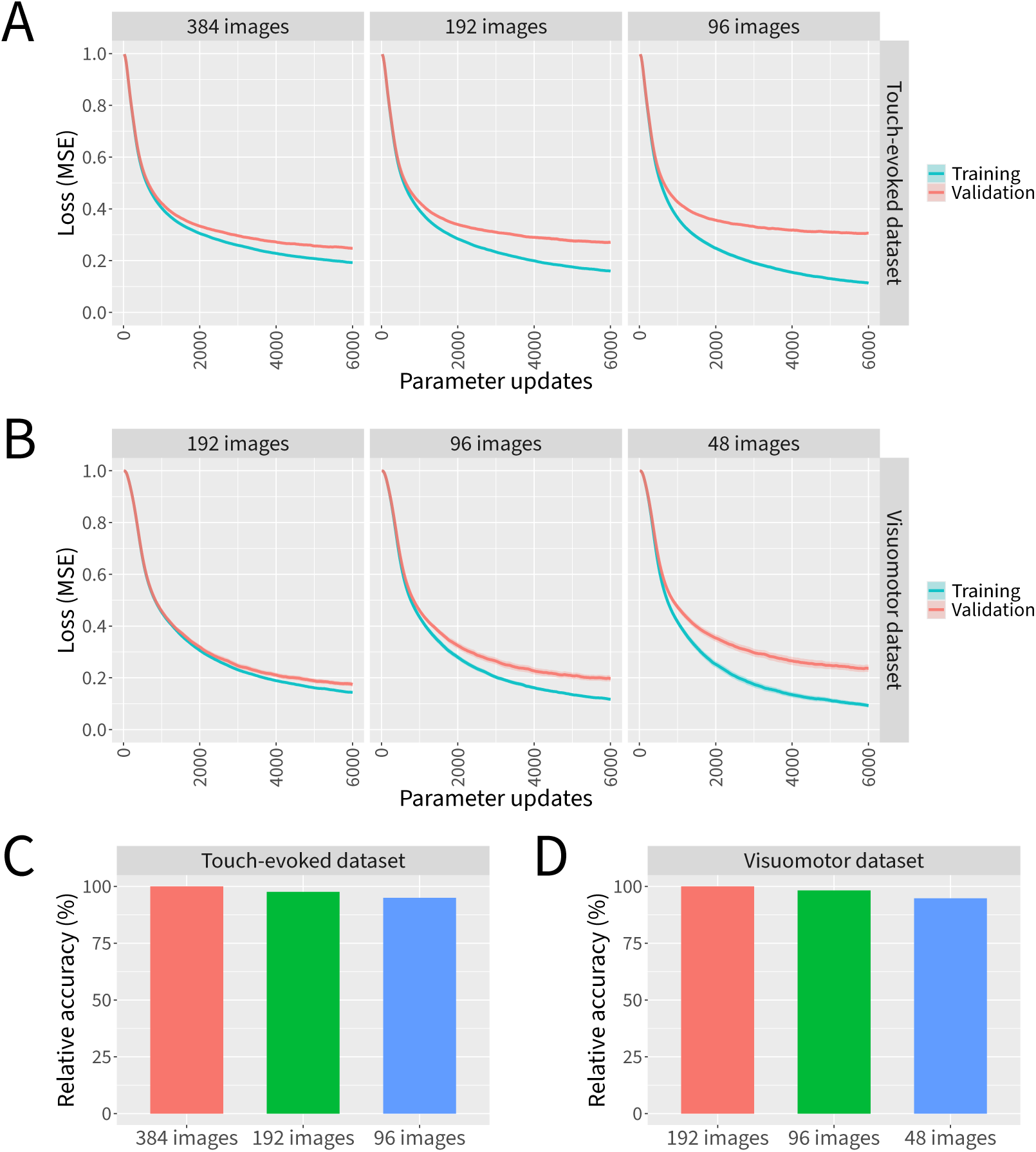
Marigold’s neural networks can be trained effectively even with limited training data. Neural networks were trained for 6000 parameter updates on increasingly ablated sets of training images while evaluating performance on a constant number of validation images. (**A**) Loss curves for the touch-evoked response dataset. (**B**) Loss curves for the visuomotor response dataset. (**C**) Relative accuracies for the touch-evoked response dataset. (**D**) Relative accuracies for the visuomotor response dataset. Data represent the mean or the mean plus and minus the standard error of the mean of 10 independent experiments in which datasets were partitioned and neural network weights were initialized using different random seeds.

Across both datasets, ablating increasing amounts of training data led to steadily decreasing training efficacy, with a disparity between the validation and training partitions in later stages of training indicating greater degrees of overfitting (Figure 6A–B). To determine to what extent this affects the accuracy of predictions, we quantified the accuracy of predictions when varying amounts of training data are available (Figure 6C–D). Because our datasets include a number of images with extreme variations in lighting conditions, occlusions, and other artifacts beyond what would normally be observed in data collected for routine biological experiments within our lab, the keypoints in some of these extreme images are not predicted well even by networks trained with all available training data. To account for this effect, we report relative accuracy as the ratio of correctly predicted keypoints for ablated datasets relative to the number of correctly predicted keypoints when all training data is available. We consider predictions to be correct when they are within 3 pixels from the ground truth label for the touch-evoked response dataset or within 1.5 pixels from the ground truth label for the visuomotor response dataset.

As data is ablated, there appears to be a steady increase in the final loss values (Figure 6A–B) and a corresponding dropoff in the number of correctly predicted keypoints (Figure 6C–D), but under the tested conditions we did not observe a catastrophic failure to converge even in the face of extremely scarce training data. We conclude that our neural networks can be trained effectively even with limited training data, particularly given that our datasets contain several orders of magnitude fewer images than many conventional computer vision datasets. However, it is likely that refinements to our data augmentation pipeline and/or adjustments to the strength of regularization techniques such as weight decay and dropout could lead to improved performance when data is more limited. Additionally, whereas we included extreme examples in our datasets to make them more challenging, better results could likely be obtained with fewer images by restricting the dataset to images from more representative behavioral recordings.

### WebAssembly implementation

Having optimized our neural network architecture using our PyTorch-based implementation and verified that it can be effectively trained with relatively little training data, we next focused on implementing this functionality within our web app. To achieve this, we implemented the necessary neural network layer and training operations in C++ and used this to generate WebAssembly, which allows such code to run within a browser at speeds approaching those of native C++ code (Haas et al., 2017; Perkel, 2024). We also explored existing neural network libraries available to web apps, but found that such libraries are generally limited in functionality compared to those available in Python and other environments. TensorFlow.js (Smilkov et al., 2019), for example, was missing implementations for several of the required layers for our chosen architecture. It also did not support gradient accumulation, which would have forced the use of smaller, less effective batch sizes to minimize our web app’s memory footprint. We also observed that many neural network implementations for the web, such as the emerging WebNN standard, are primarily focused on inference and lack support for training.

To evaluate the effectiveness of our WebAssembly-based approach, we compared the resulting training and inference speeds to those of our PyTorch implementation (Figure 7). In terms of training speed, Marigold slightly to moderately outperformed PyTorch across neural networks configured for both the touch-evoked response and visuomotor response datasets. In terms of inference speed, Marigold slightly outperformed PyTorch for neural networks configured for the visuomotor response dataset, but was moderately outperformed by PyTorch for neural networks configured for the touch-evoked response dataset. Overall, our implementation achieves speeds that are competitive on CPU with the highly optimized PyTorch library. This is particularly noteworthy given the relatively limited access to memory and CPU features that web apps are permitted.

**Figure 7.**
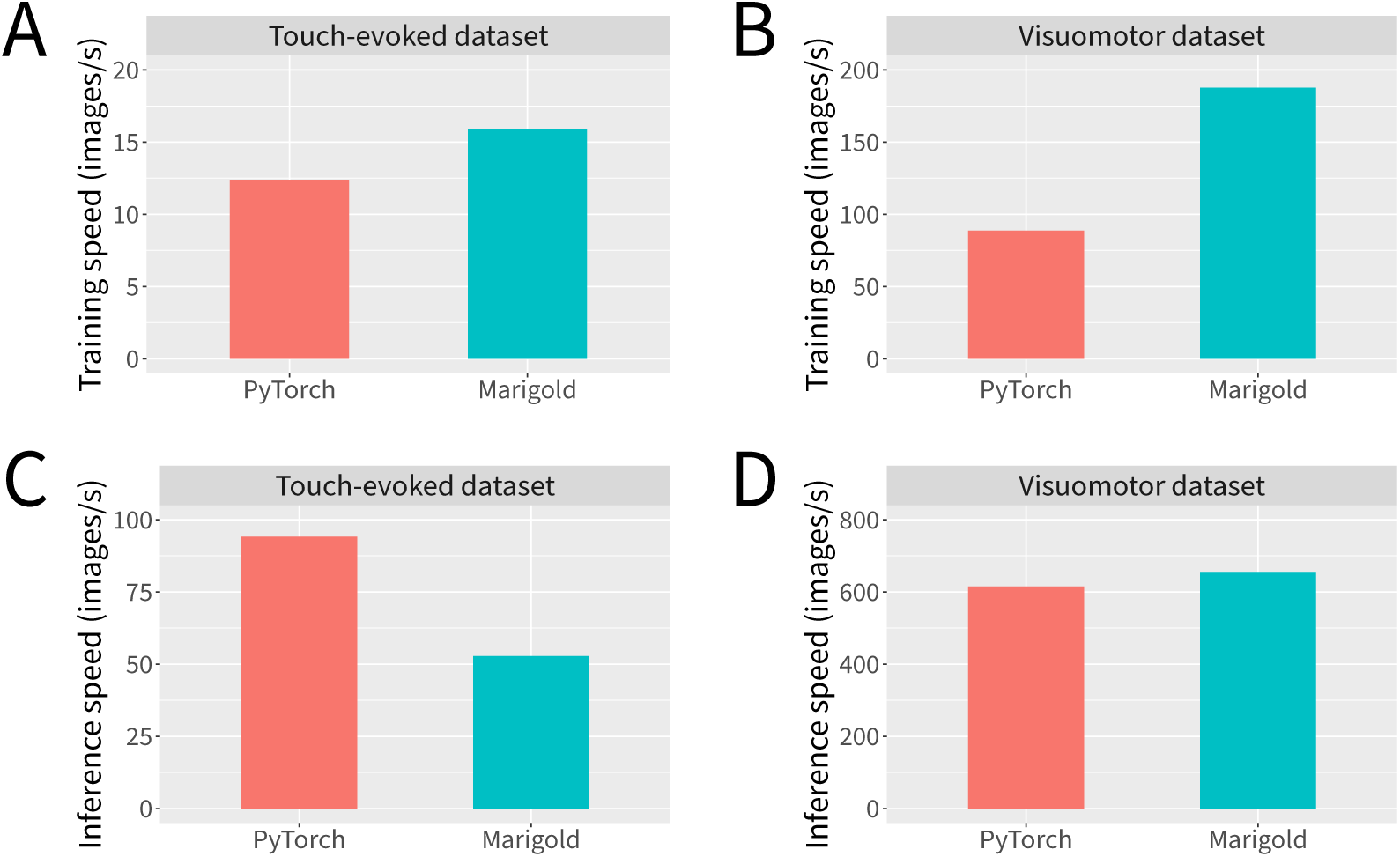
Marigold’s WebAssembly-based neural network implementation achieves CPU performance competitive with that of PyTorch. (**A**) Training speeds for the touch-evoked response dataset. (**B**) Training speeds for the visuomotor response dataset. (**C**) Inference speeds for the touch-evoked response dataset. (**D**) Inference speeds for the visuomotor response dataset. Measurements reflect single-threaded CPU performance. Training speeds were measured using gradient accumulation with a batch size of 1 to obtain an effective batch size of 16. Inference speeds were measured with a batch size of 1. Data represent the mean of 10 independent measurements.

### Effects of genotype and touch location in *slc1a2b^tk57/tk57^* and wild-type sibling touch-evoked response behavior

Using Marigold, we investigated novel aspects of the *techno trousers* (*tnt*) locomotor phenotype (Granato et al., 1996; McKeown et al., 2012). This mutant, henceforth referred to as *slc1a2b^tk57/tk57^*, harbors a loss-of-function missense mutation in *slc1a2b* (Ensembl ID: ENSDARG00000102453) located on chromosome 25, which encodes Eaat2b, a glutamate transporter predominantly expressed in astroglia (Hotz et al., 2022; McKeown et al., 2012). We note that this gene and its protein product have also been referred to as *slc1a2a* and Eaat2a, respectively (Gesemann et al., 2010; Hotz et al., 2022); however, current genome databases indicate that these particular identifiers refer to a distinct gene found on chromosome 7 (Ensembl ID: ENSDARG00000052138), and its protein product (Breuer et al., 2019). *slc1a2b^tk57/tk57^* embryos demonstrate a hyperactive response to touch by 48 hpf; however, the larvae are effectively paralyzed and shorter along the rostral–caudal axis by 96 hpf (McKeown et al., 2012). Small twitch responses to touch can be observed at 96 hpf, suggesting the lack of response is due to motor issues rather than sensory issues. Because the auditory and visual systems are not fully developed by 96 hpf, tactile stimulation at earlier time points is the most robust way to elicit this hyperactive phenotype.

Reticulospinal neurons, such as the Mauthner cell and its homologs (collectively referred to as the Mauthner array), are the primary mediators of the short latency touch-evoked response. Head-stimulated responses recruit the full Mauthner array while tail-stimulated responses recruit the Mauthner cell alone, resulting in kinematically distinct responses based upon touch location (K. S. Liu & Fetcho, 1999). By comparing touch-evoked responses of head- and tail-stimulated *slc1a2b^tk57/tk57^* embryos, we reasoned that we could uncover novel aspects of the *slc1a2b^tk57/tk57^* locomotor phenotype.

Following head stimulation, wild-type embryos reliably exhibit a high amplitude body bend, reorienting the embryo, followed by lower amplitude undulations, allowing the embryo to swim away from the perceived threat (Figure 8A). Head-stimulated *slc1a2b^tk57/tk57^* embryos swim significantly longer and farther than wild-type controls (two-way ANOVA, F(1,86) = 59.58, p < 0.001 and F(1,86) = 57.81, p < 0.001 for duration and distance, respectively) (Figure 8B–C). Consistent with previous literature (McKeown et al., 2012), *slc1a2b^tk57/tk57^* embryos perform more high amplitude body bends during the escape response (two-way ANOVA, F(1,86) = 35.88, p < 0.001) (Figure 8D). However, the number of high amplitude body bends performed across head- and tail-stimulated embryos did not vary by genotype (two-way ANOVA, F(1,86) = 0.00, p >= 0.05). The distribution of high amplitude body bends throughout the startle response differs markedly between *slc1a2b^tk57/tk57^* and wild-type embryos. *slc1a2b^tk57/tk57^* embryos perform extended stretches of high amplitude body bends throughout the responses (Figure 8E; Figure 9A). Of all embryos that perform a high amplitude body bend, the distribution of *slc1a2b^tk57/tk57^* high amplitude body bends features more body bends occurring later in the response (Figure 9A–B). We further investigated characteristics of the first body bend within each startle response. Regardless of genotype, the initial body bend of tail-stimulated embryos is significantly lower in amplitude than that of head-stimulated embryos, often not crossing the 110° threshold (two-way ANOVA, F(1,86) = 79.703, p < 0.001) (Figure 9C). Tail-stimulated embryos also reached their maximum angle significantly earlier in the response (two-way ANOVA, F(1,86) = 44.833, p < 0.001) (Figure 9D). Interestingly, we note genotype- and touch location-dependent effects on the maximum body angle of the startle response. Among wild-type embryos, the maximum body bend had a significantly lower amplitude in tail-stimulated embryos than in head-stimulated embryos (two-way ANOVA, F(1,86) = 36.78, p < 0.001). In contrast, the maximum angle for tail-stimulated *slc1a2b^tk57/tk57^* responses is nearly indistinguishable from head-stimulated *slc1a2b^tk57/tk57^* responses (two-way ANOVA interaction, F(1,86) = 14.45, p < 0.001) (Figure 9E). To determine whether embryos execute the maximum angle after the initial body bend, we measured the time required for the embryo to reach the maximum body angle of the response. *slc1a2b^tk57/tk57^* embryos perform their maximum body bend significantly later in the response than wild-type embryos (two-way ANOVA, F(1,86) = 25.26, p < 0.001) (Figure 9F).

**Figure 8.**
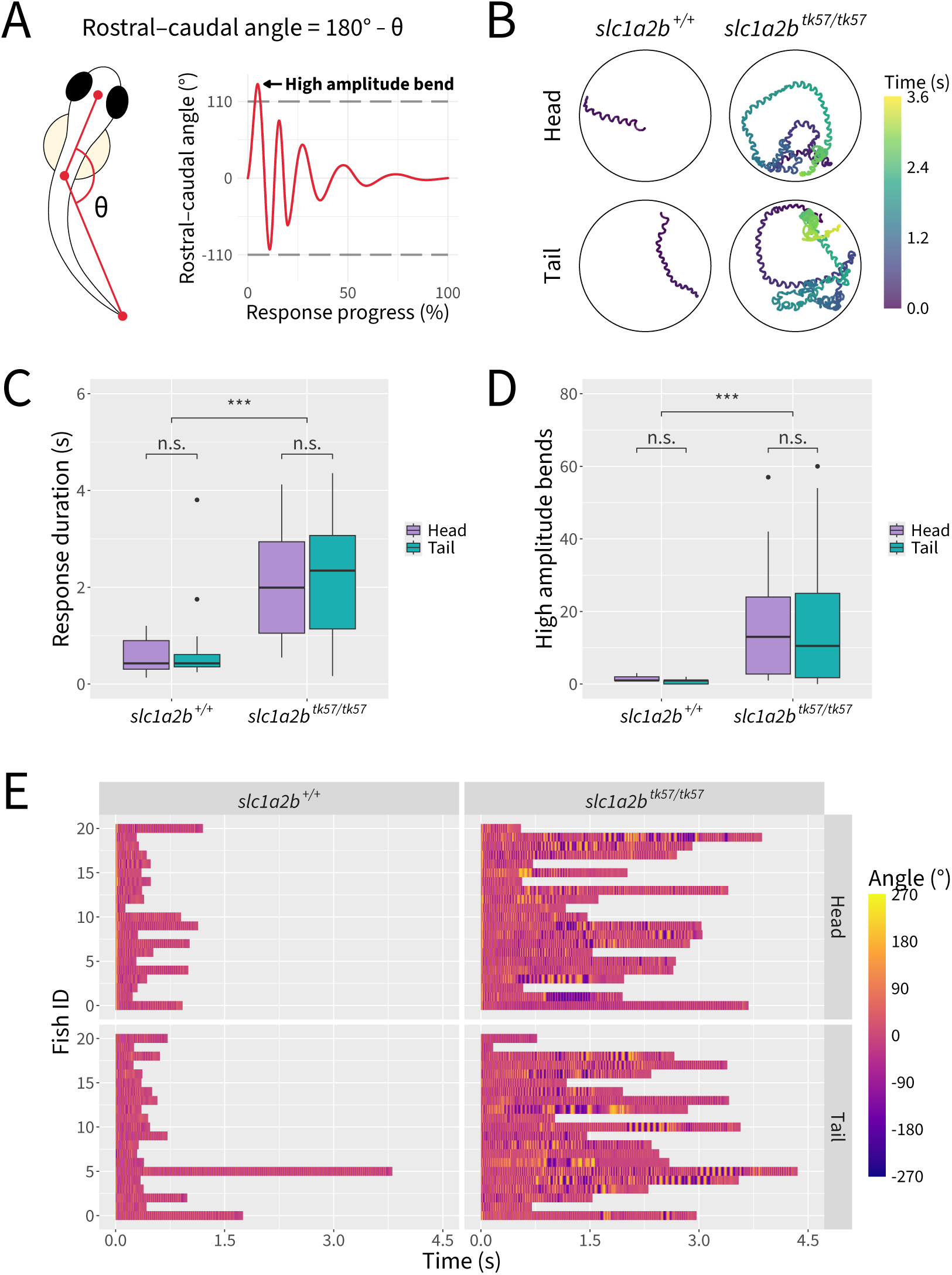
*slc1a2b^tk57/tk57^* mutant zebrafish exhibit hyperactive touch-evoked startle behavior regardless of the touch location. (**A**) Schematic illustrating calculation of rostral–caudal angle and quantification of high amplitude body bends. (**B**) Representative trajectory traces. (**C**) Quantification of response durations. (**D**) Quantification of high amplitude body bends. (**E**) Visualization of individual rostral–caudal angles over time, with three fish omitted at random in order to show equal numbers of fish across conditions. Statistical significance determined by two-way ANOVA with Tukey post hoc; n.s.: not significant; ***: p < 0.001; n = 21–24 fish per condition across 3 independent experiments.

**Figure 9.**
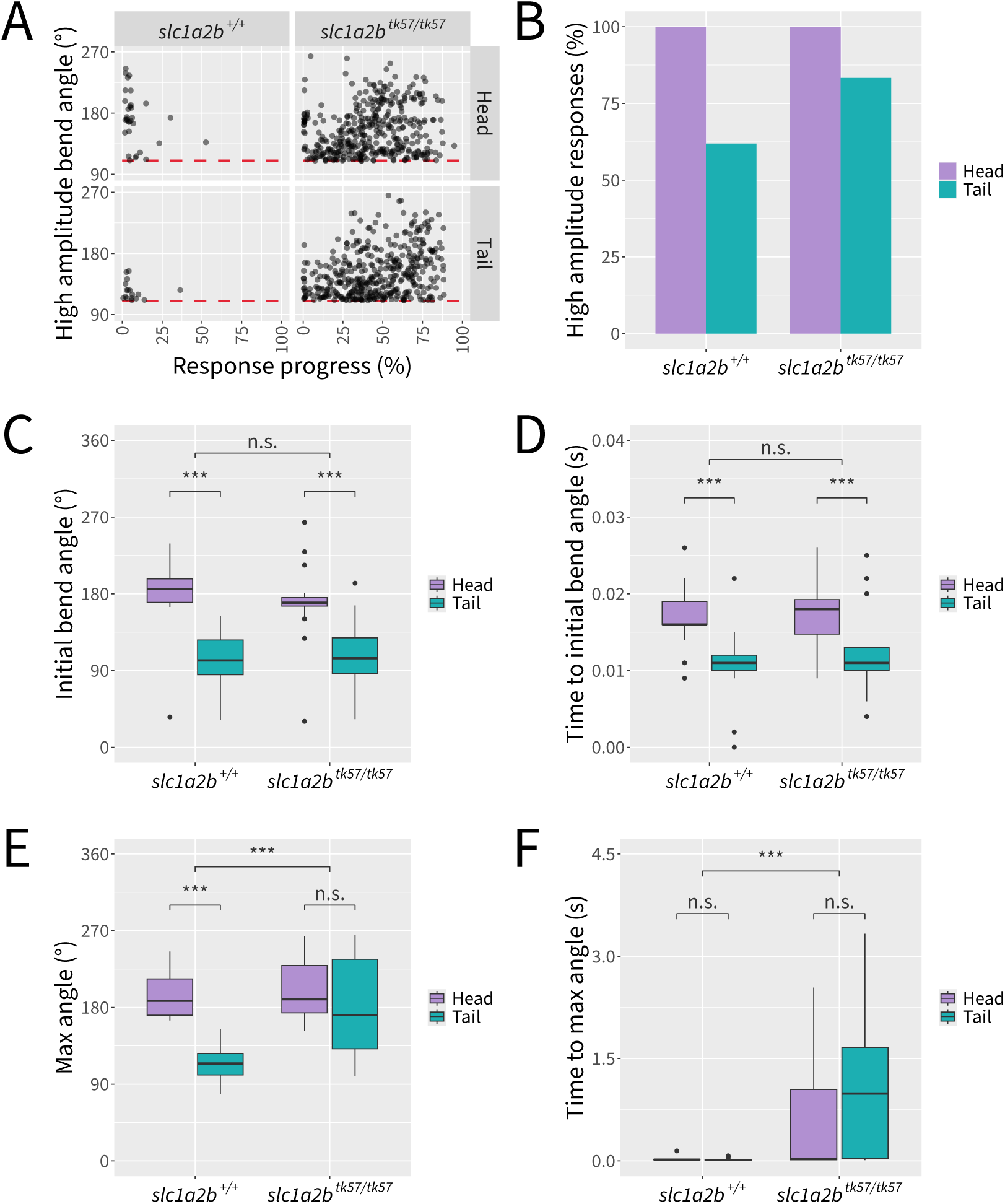
*slc1a2b^tk57/tk57^* and wild-type embryos perform rostral–caudal body bends dependent on interactions between genotype and touch location. (**A**) Visualization of absolute rostral–caudal angle magnitudes at the peaks of high amplitude body bends, depicted over time according to overall response progress. Three fish were omitted at random in order to represent equal numbers of fish across conditions. Red line indicates the threshold of 110° for classification as a high amplitude body bend. (**B**) Percentage of responses which contain one or more high amplitude body bends. (**C**) Quantification of peak angle of first body bend. (**D**) Quantification of time to peak angle of first body bend. (**E**) Quantification of maximum angle across the entire response. (**F**) Quantification of time to maximum angle across the entire response. Statistical significance determined by two-way ANOVA with Tukey post hoc; n.s.: not significant; ***: p < 0.001; n = 21–24 fish per condition across 3 independent experiments.

### Effects of developmental stage and feeding on larval visuomotor response behavior

To demonstrate additional capabilities of Marigold, particularly its ability to accurately and efficiently analyze the behavior of larval zebrafish in multiwell plates, we examined the effects of developmental stage and feeding on the larval visuomotor response (Brockerhoff et al., 1997; Burgess & Granato, 2007a; Lee et al., 2019; Randlett et al., 2019). Larvae were fed or not fed once daily from 5 dpf through 7 dpf and behavior was recorded in 24-well plates 4–6 h after feeding at 5 dpf and 7 dpf. To examine the effects of age and feeding across multiple behavioral modes, we recorded fish during periods of dark adaptation, light stimulus, light adaptation, and dark stimulus (Figure 10A).

**Figure 10.**
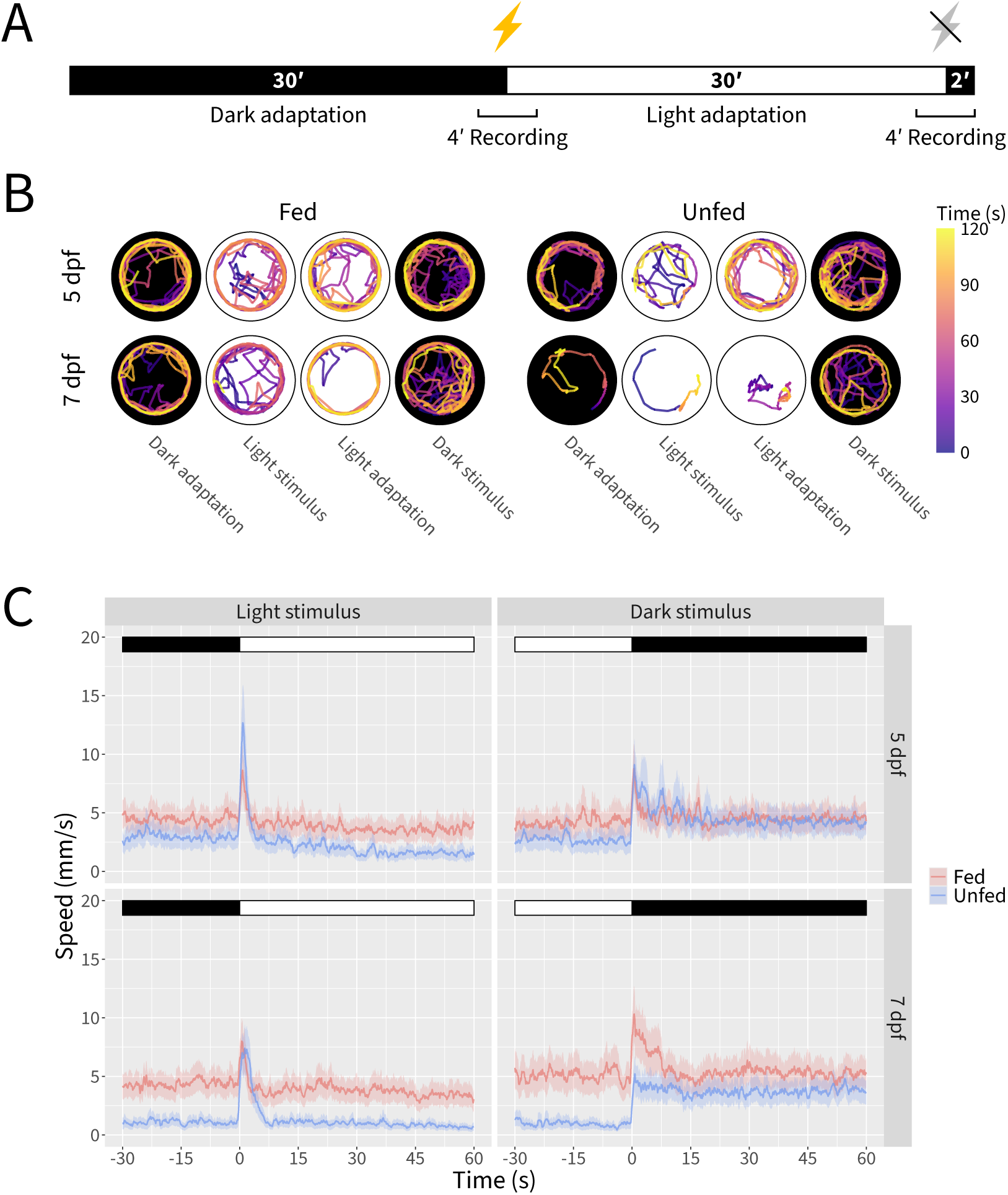
Visuomotor response trajectory traces and speed plots illustrate reduced swimming in unfed larvae. (**A**) Schematic of visuomotor response recording paradigm. (**B**) Representative trajectory traces for fed and unfed fish at 5 and 7 dpf during periods of dark adaptation, light stimulus, light adaptation, and dark stimulus. (**C**) Speed plots for fed and unfed fish at 5 and 7 dpf during periods of dark adaptation, light stimulus, light adaptation, and dark stimulus. Traces represent the mean plus and minus the standard error of the mean after applying a mean filter with a sliding window corresponding to 1 s. Data represents n = 44–46 fish per condition across 3 independent experiments.

Visualization of fish trajectories during the four recording periods suggested reduced swimming in unfed fish, particularly at 7 dpf (Figure 10B). Smoothed mean speed traces provided further insight into this pattern (Figure 10C). During dark adaptation, both fed and unfed fish maintained stable activity levels, however this baseline activity level was lower in unfed fish, with the effect becoming more pronounced at 7 dpf. Interestingly, the baseline activity level was essentially identical across fed fish regardless of age. Immediately following the light stimulus, both fed and unfed fish mobilized a rapid burst of activity before returning to the baseline activity level. Fed and unfed fish exhibited similar level of activity during light adaptation and dark adaptation, with unfed fish again showing a lower level of activity which was more pronounced at 7 dpf. Strikingly, whereas the acute reaction to the dark stimulus was followed by a return to baseline in fed fish (albeit more gradually than was observed following the light stimulus), unfed fish maintained a heightened level of activity which resembled the baseline activity level of fed fish.

Analysis of maximum speed levels during the 5 seconds immediately following light or dark stimuli revealed no significant differences or interactions between age and feeding status for light stimuli (two-way ANOVA, F(1,174) = 3.537 for age, 1.608 for diet, and 2.401 for interaction, p >= 0.05 in all three cases). In contrast, immediately following dark stimuli unfed fish exhibited an attenuated response at 7 dpf, but not at 5 dpf (Figure 10C; Figure 11A), reflecting a main effect of diet (two-way ANOVA, F(1,174) = 21.659, p < 0.001) and an interaction between age and diet (two-way ANOVA, F(1,174) = 29.881, p < 0.001), but no main effect of age (two-way ANOVA, F(1,174) = 0.107, p >= 0.05). Further insights were gleaned by examining the number of swim bouts, which we define as beginning when a fish’s speed exceeds 2 mm/s and ending when the fish’s speed drops below this threshold. Here, a main effect of diet was consistently observed during periods of dark adaptation, light stimulus, light adaptation, and dark stimulus (two-way ANOVA, F(1,178) = 144.94, 100.513, 48.606, and 27.127, respectively, p < 0.001 during all four periods). However, a significant main effect of age was observed during only the first three of these periods (Figure 11B) (two-way ANOVA, F(1,178) = 23.61, 11.855, 9.129, and 3.535, respectively, p < 0.001, p < 0.001, p < 0.01, and p >= 0.05, respectively). The interaction between age and diet followed a similar pattern (Figure 11B) (two-way ANOVA, F(1,178) = 26.08, 9.942, 8.146, and 0.504, respectively, p < 0.001, p < 0.01, p < 0.01, and p >= 0.05, respectively). In addition to the number of swim bouts, we also qualitatively examined the vigor of swim bouts by visualizing the durations and peak speeds of swim bouts for each fish (Figure 11C). Whereas the swim bouts of fed and unfed fish generally align at 5 dpf, by 7 dpf unfed fish exhibit less vigorous swimming behavior compared to fed fish as evidenced by their shorter and lower-speed swim bouts.

**Figure 11.**
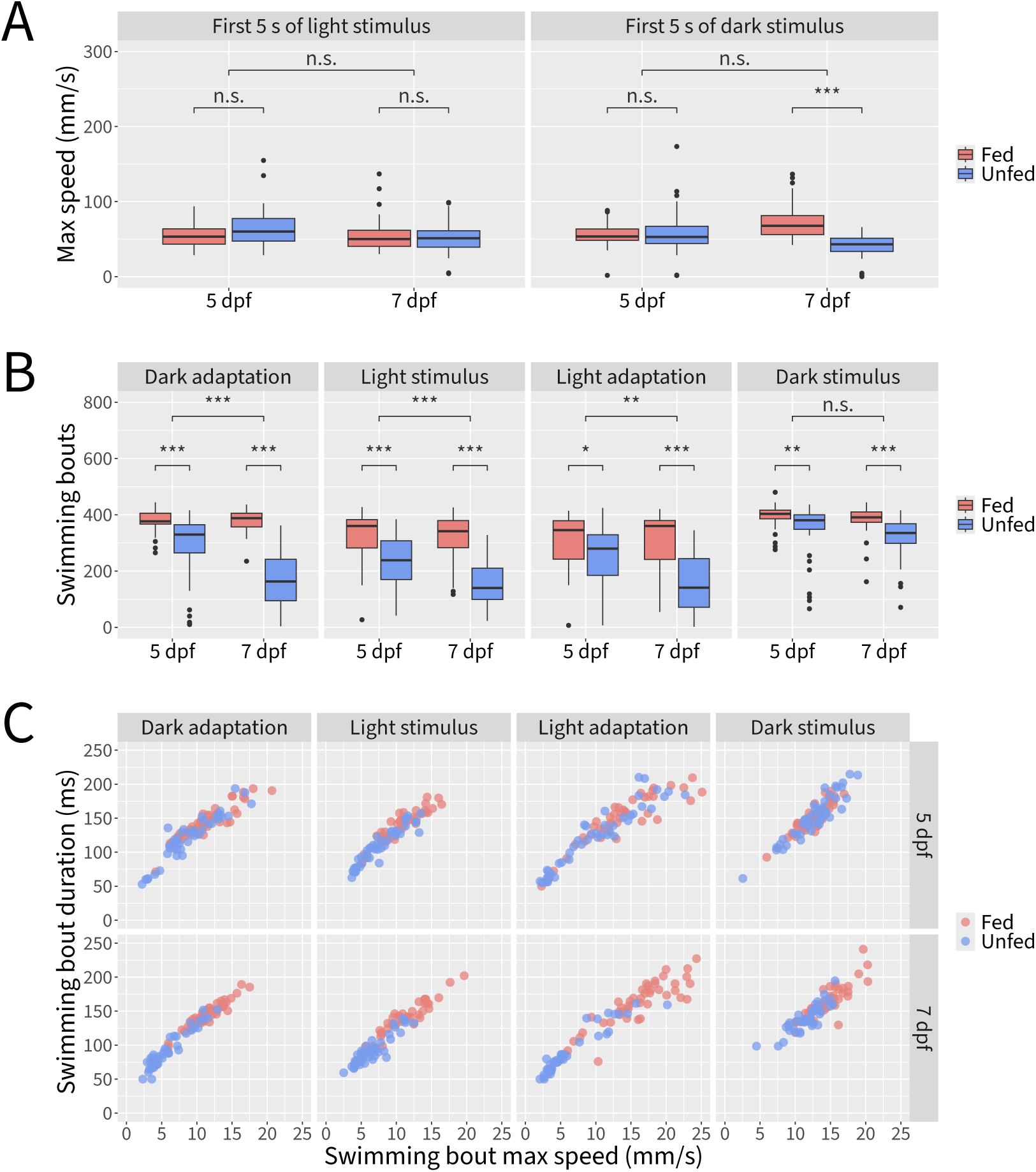
Detailed behavioral profiling reveals effects and interactions of age and diet on larval visuomotor responses. (**A**) Quantification of maximum speed during the first 5 s of the response to light and dark stimuli for fed and unfed fish at 5 and 7 dpf. (**B**) Quantification of swimming bouts for fed and unfed fish at 5 and 7 dpf during periods of dark adaptation, light stimulus, light adaptation, and dark stimulus. (**C**) Visualization of mean swimming bout duration and mean swimming bout maximum speed for fed and unfed fish at 5 and 7 dpf during periods of dark adaptation, light stimulus, light adaptation, and dark stimulus. Statistical significance determined by two-way ANOVA with Tukey post hoc; n.s.: not significant; *: p < 0.05; **: p < 0.01; ***: p < 0.001; n = 44–46 fish per condition across 3 independent experiments.

## Discussion

### Biological experiments

To demonstrate the utility of Marigold, we analyzed touch-evoked responses in *slc1a2b^tk57/tk57^* embryos and their wild type siblings, with the stimulus being applied to either the head or tail. We first replicated the known phenotypic differences in the touch-evoked response between *slc1a2b^tk57/tk57^* embryos and their wild type siblings when applying the stimulus to the head, including increases in response duration and the number of high amplitude body bends in *slc1a2b^tk57/tk57^* embryos. We then explored whether applying a stimulus to either the head or tail would further reveal distinct kinematic profiles in *slc1a2b^tk57/tk57^* embryos. Within tail-stimulated responses, fewer embryos demonstrate an initial body bend that passes the high amplitude threshold, compared to head-stimulated embryos, regardless of genotype. Because the embryo is already oriented away from the perceived threat when touched on the tail, reorientation via a high amplitude body bend may be less advantageous in this scenario.

We further show that while there are stimulus location differences in the maximum rostral– caudal angle that a wild-type embryo performs, those differences are lost in *slc1a2b^tk57/tk57^* embryos. Both head- and tail-stimulated *slc1a2b^tk57/tk57^* embryos reach the maximum rostral– caudal angle later in the response than wild-type embryos, which could help explain why no touch location differences were found for the maximum angle of *slc1a2b^tk57/tk57^* responses. Although *slc1a2b* is not expressed in the Mauthner cell, *slc1a2b*-expressing glial cells surround the Mauthner array at 48 hpf (Hotz et al., 2022; McKeown et al., 2012). Based on the data presented here, we propose that the disrupted kinematic profile of *slc1a2b^tk57/tk57^* embryos may be due to a reactivation of escape circuitry late into the response.

Interestingly, *slc1a2b* mutants harboring nonsense mutations are able to swim spontaneously at 5 dpf (Hotz et al., 2022), whereas the *slc1a2b^tk57/tk57^* loss-of-function missense allele described here and in McKeown et al., 2012 is fully paralyzed by 96 hpf. This difference could be explained by the nature of the mutations; premature termination codons generated using CRISPR-Cas9 have been shown to trigger genetic compensation via transcriptional adaptation (El-Brolosy et al., 2019; Sztal & Stainier, 2020). Additionally, the *slc1a2b^tk57/tk57^* allele described here is maintained on a TL background, while Hotz et al., 2022 report using WIK and Tübingen wild-type strains. As notable differences have been reported between wild-type zebrafish strains (Audira et al., 2020; Maximino et al., 2013), this could also explain the difference in phenotypic severity. Lastly, Hotz et al., 2022 describe a mutation in exon 3 of *slc1a2b* while the mutation underlying *slc1a2b^tk57/tk57^* is located in exon 7 of *slc1a2b* (McKeown et al., 2012).

We also used Marigold to examine visuomotor responses in wild-type larvae at two developmental stages and in fed and unfed states. Larval responses to light differ across developmental time (de Esch et al., 2012; Y. Liu et al., 2015). The first startle responses to visual stimulation can be observed around 3 dpf (Easter & Nicola, 1996). By 5 dpf, larvae inflate their swim bladders, exhibit robust swimming behavior, and actively hunt for food. Larvae utilize nutrients from their yolks until at least 7 dpf, and it has been shown that feeding can be delayed until 8 dpf without negatively affecting juvenile survival or growth (Hernandez et al., 2018; Lucore & Connaughton, 2021; A. V. Schwartz et al., 2021). A previous study examining the effects of feeding on larval swim behavior, visual stimuli avoidance, and inter-fish distance reported significant behavioral differences between fed and unfed fish at 6 and 7 dpf, but no significant effects at 5 dpf (Clift et al., 2014). This observation led to the recommendation that 5 dpf generally be preferred for behavioral experiments. However, the effect of feeding has not previously been evaluated with respect to visuomotor responses.

While our data confirm that feeding status has a more pronounced effect on visuomotor response behavior at 7 dpf than at 5 dpf, we also provide evidence for significant effects of feeding on behavioral parameters even at 5 dpf (Brockerhoff et al., 1997; Burgess & Granato, 2007a; Lee et al., 2019; Randlett et al., 2019). Thus, our results highlight the importance of reporting and controlling for developmental stage and feeding status in visuomotor response assays.

In both sets of experiments, the in-depth behavioral profiling facilitated by Marigold led to novel findings, despite the *slc1a2b^tk57/tk57^* mutant having been previously characterized and visuomotor responses being a widely adopted experimental paradigm.

### Neural network experiments

To minimize the processing and memory demands of in-browser, on-device neural networks and thereby make them feasible for our zebrafish pose tracking web app, we introduced new neural network variants originating from a series of macro-architectural and micro-architectural design decisions.

Our switch from a hierarchical architecture to an isotropic architecture led to considerably faster training and inference speeds and a reduced memory footprint. Isotropic (also sometimes called isometric) architectures are relatively unexplored. Sandler, Howard, et al., 2019 introduced a form of isometric neural networks and noted its memory efficiency and other intriguing properties, although they ultimately found that it did not perform as well for image classification as more traditional hierarchical architectures. More recently, the Vision Transformer family of architectures, which draw inspiration from transformer-based architectures for natural language processing, used isotropic design as a way to reduce the computational cost of the self-attention mechanism used in transformers (Dosovitskiy et al., 2021; Vaswani et al., 2017). Some works have been successful in modifying Vision Transformer-like architectures to more closely resemble the hierarchical architectures traditionally used for computer vision (Z. Liu et al., 2022; Yu et al., 2022). Meanwhile, other works have explored whether the patch-based representation used by isotropic architectures may be useful in and of itself, perhaps underlying much of the success of Vision Transformers (Feng et al., 2022; Tolstikhin et al., 2021; Trockman & Kolter, 2022).

Our switch from Batch Normalization to Instance Normalization led to improved training dynamics and allowed us to use gradient accumulation to reduce memory requirements. Although there has been a strong preference for Batch Normalization in neural networks for computer vision, alternatives such as Layer, Group, and Instance Normalization have increasingly found niches in which they are effective. Isensee et al., 2021, for example, find Instance Normalization useful in U-Net based architectures for semantic segmentation of biomedical images. Also using U-Net for biomedical image segmentation, Neubig and Kist, 2023 use neural architecture search to explore the performance of different normalization methods on a block-by-block basis, finding that Instance Normalization is preferred at most, but, interestingly, not all, positions. The ConvNeXt family of neural networks, which draws inspiration from Vision Transformer-like architectures where Layer Normalization is predominant, found a benefit to switching from Batch Normalization to Layer Normalization, but did not test other normalization methods (Z. Liu et al., 2022; Woo et al., 2023). Another line of work, which we did not pursue, seeks to eliminate normalization layers altogether (e.g., Brock et al., 2021; Gadhikar and Burkholz, 2022; Zhang et al., 2019; Zhang et al., 2022).

We also achieved greater training and inference speeds by removing some of the normalization and activation layers present in the MobileNetV3 block. That the choice of which normalization and activation layers to retain led to different training outcomes emphasizes the importance of being strategic in the adoption of this approach. Z. Liu et al., 2022 also find some benefit to using fewer normalization and activation layers, although they use a standard ResNet bottleneck as their starting point and arrive at a somewhat different final configuration of layers.

As the state of the art in machine learning continues to demand ever-increasing computational resources to power deep neural network training and inference for computer vision and natural language processing applications, the associated carbon footprint exerts an increasingly negative impact on the environment (R. Schwartz et al., 2019; Strubell et al., 2019; Verdecchia et al., 2023; Xu et al., 2023). Thus, we feel the machine learning community has a moral responsibility to design more efficient neural networks. Our design choices highlight ways in which architectural optimizations can help to minimize the processing and memory demands of neural networks while leading to faster training convergence and inference speed. It is perhaps worth noting that we achieved our goal of reducing the processing and memory demands of neural networks for animal pose estimation primarily through architectural improvements, but a number of promising alternative strategies could have been explored instead or in addition, such as knowledge distillation, quantization, and pruning (Dantas et al., 2024).

### Comparison to existing free and open source software

Numerous free and open source software solutions are available for zebrafish behavioral analysis, encompassing a wide range of tracking capabilities, implementation details, hardware requirements, expectations for programming experience, and degrees of specificity to zebrafish. While a comprehensive review of all such software is beyond the scope of this work, we summarize a number of representative tools below.

Among animal-agnostic solutions, several recently introduced tools have sought to make advances in machine learning-based pose estimation techniques available to researchers interested in analyzing animal behavior (Goodwin et al., 2024; Graving et al., 2019; Mathis et al., 2018; Pereira et al., 2019). These tools have tremendously improved the precision with which animal pose tracking can be performed, but are not without limitations. Because these tools commonly use off-the-shelf neural network architectures which were originally designed for other datasets or tasks, the resulting models are frequently overparameterized and their training can be inefficient, often requiring the use of expensive GPUs. Additionally, such software can be challenging to install due to complex dependencies on language runtimes, libraries, and device drivers, as well as difficult to use due to workflows requiring integration of GUI-, command line-, and cloud-based elements. Additionally, most of these tools lack support for tracking animals in multi-well plates, a popular experimental paradigm for high-throughput behavioral studies using larval zebrafish.

Many zebrafish-specific or small animal-specific tools have also been introduced. Relatively few of these use machine learning, with most instead relying on traditional computer vision techniques such as background subtraction, centroid tracking, and following local pixel intensity cues to identify different parts of the fish (Audira et al., 2018; Burgess & Granato, 2007b; Chen et al., 2023; Clift et al., 2014; Joo et al., 2020; Marques et al., 2018; Mirat et al., 2013). These approaches can be adequate, but are often brittle, requiring carefully controlled experimental setups to deliver reliable results and breaking down under adverse conditions such as the introduction of foreign objects into the field of view. Additionally, some of these tools are limited to only tracking a single keypoint. Those tools that incorporate machine learning-based tracking typically do so by drawing on one of the more general purpose tools described above, such as DeepLabCut, to provide the required functionality, and thus can inherit many of the same limitations as these tools such as a complex installation process and the requirement for a powerful GPU (Gore et al., 2023; Guilbeault et al., 2021; Štih et al., 2019; Tadres & Louis, 2020; Zhu et al., 2023). A number of tools combine traditional or machine learning-based tracking abilities with camera and other hardware integration, including affordable hardware setups such as those based on Raspberry Pi, to drive data collection and stimulus delivery for experiments (Guilbeault et al., 2021; Joo et al., 2020; Singh et al., 2023; Štih et al., 2019; Tadres & Louis, 2020).

Marigold represents a notable departure from existing software primarily in its relative lack of hardware requirements, elimination of installation procedures, and focus on making robust, machine learning-based pose tracking more widely accessible to both students and researchers. To our knowledge, Marigold is the first free and open source software to perform machine learning-based animal pose tracking for behavioral analysis entirely in-browser and on-device, an approach which allows us to meaningfully lower the financial and technical barrier for performing this type of analysis. By implementing our software as a web app, we are able to take advantage of the web platform to deliver a rich user interface supporting a highly streamlined workflow, all without requiring any installation. The web app implementation also allows Marigold to run entirely within the security sandbox provided by the user’s web browser. Moreover, by designing a neural network architecture from the ground up to perform highly efficient animal pose estimation and implementing this architecture using the recently introduced Web-Assembly technology (Haas et al., 2017; Perkel, 2024), we are able to achieve reasonable training and inference speeds even on modestly powered computers lacking a dedicated GPU and despite the limited access to memory and computational resources web apps are permitted.

### Future directions

Marigold’s ability to allow the user to create their own dataset and train custom models, while maintaining a focus on functionality that is of particular interest to the zebrafish research community, makes the program applicable to a broad range of zebrafish behavioral experiments. For instance, in addition to the applications demonstrated here (i.e., analysis of touch-evoked and visuomotor responses), we also have successfully used Marigold to analyze spinalized larvae, head-embedded larvae, and adult fish (data not shown). However, Marigold is currently limited to analyzing a single animal in each region of interest (ROI), a decision made to maintain the program’s simplicity and our focus on embryonic and early larval stages of zebrafish development, when social interaction is minimal (Stednitz & Washbourne, 2020). Future work could extend Marigold to support the analysis of multiple fish in each ROI and for three-dimensional pose tracking, since both social interaction and three-dimensional swimming become more prominent during late larval and juvenile development (Macrì et al., 2017; Stednitz & Washbourne, 2020). Additionally, while we have successfully used Marigold for pose tracking of other animal species such as mice (data not shown), there is clearly an opportunity to more formally extend the web app-based approach of Marigold for pose tracking in other species and to adapt it for other types of behavioral analysis.

Finally, our WebAssembly-based implementation was able to obtain reasonable performance on CPU, making our web app compatible with a wide range of platforms and devices and making it particularly valuable in low-resource settings. However, for devices equipped with a discrete GPU Marigold may leave performance on the table, especially when it comes to larger neural networks such as those we trained for our touch-evoked response dataset. The emerging WebGPU standard could provide a means for Marigold to take advantage of advanced GPUs when they are available, helping to close this performance gap. In the meantime, we note that Marigold may not be suited for all types of behavioral recordings. Marigold is quite efficient for high-throughput, low-resolution tracking with few keypoints, as exemplified by the models trained for analyzing visuomotor responses. It is also fast enough for high-resolution tracking with many keypoints when the recordings are relatively short, as exemplified by the models trained for analyzing touch-evoked responses, but may not be fast enough to make such high-resolution, many-keypoint tracking feasible for extended recordings.

## Conclusions

Much of the existing software tools used for zebrafish behavioral analysis have constraints that limit their utility, including being cost prohibitive or requiring substantial programming experience. Here, we describe Marigold, which aims to address these limitations. It is a free and open source web app that utilizes machine learning to perform zebrafish pose tracking. Marigold features efficient neural networks that allow for reasonable training and inference speeds even on basic laptop computers. We demonstrate the utility of Marigold by using it to uncover novel aspects of the *tnt*mutant touch-evoked escape response and of the effects of developmental stage and feeding on the larval visuomotor response. We expect that Marigold will serve as a user-friendly tool to aid the zebrafish community in conducting robust, high-throughput behavioral analysis.

## Acknowledgments

We thank Hanna E. Dorman Barclay and Caroline Martin for excellent fish care. We thank all members of the Downes Lab, as well as the University of Massachusetts Amherst and Amherst College zebrafish communities, for thoughtful discussion. We are extremely grateful to Scott Alfeld for helpful discussion and feedback on an early version of the manuscript.

## Author contributions

GT, JGT, and GBD acquired funding for the project. GT, RMR, WB, JGT, and GBD conceptualized the software and the biological experiments. GT conceptualized and conducted the computational experiments and developed the software. GT and GM collected the training data and conducted the biological experiments. RMR and BEC labeled the training data. GT, RMR, and WB analyzed the biological experiments and prepared the figures. GT, RMR, WB, BEC, GBD, and JGT wrote the manuscript. All authors read and approved the final manuscript.

## Data availability

The datasets used and/or analyzed during this study are available from the corresponding authors upon reasonable request. The source code for Marigold is licensed under the GNU General Public License version 3 (GPLv3) and is available at: https://github.com/downeslab/marigold. The Marigold web app itself is available at: https://downeslab.github.io/marigold/.

