## Supplementary Figures for "Marigold: A machine learning-based web app for zebrafish pose tracking"

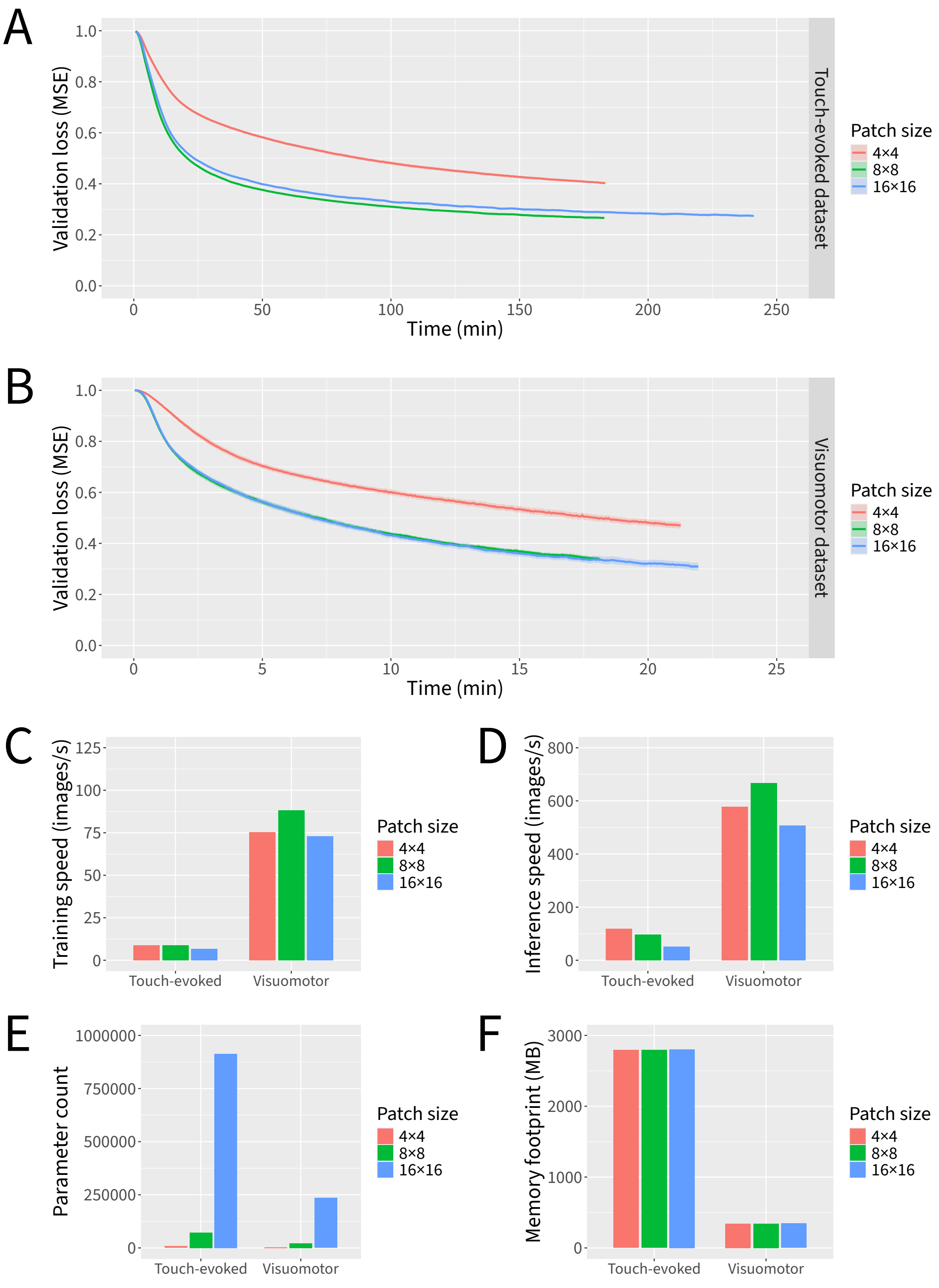


**Supplementary Figure 1. Ablation experiment for the macro-architectural design step.** Neural networks were trained for 6000 parameter updates for each dataset and patch size. (**A**) Loss curves for the touch-evoked response dataset. (**B**) Loss curves for the visuomotor response dataset. (**C**) Training speeds. (**D**) Inference speeds. (**E**) Parameter counts. (**F**) Theoretical minimum memory footprints. Data represent the mean plus and minus the standard error of the mean of 10 independent experiments in which datasets were partitioned and neural network weights were initialized using different random seeds.

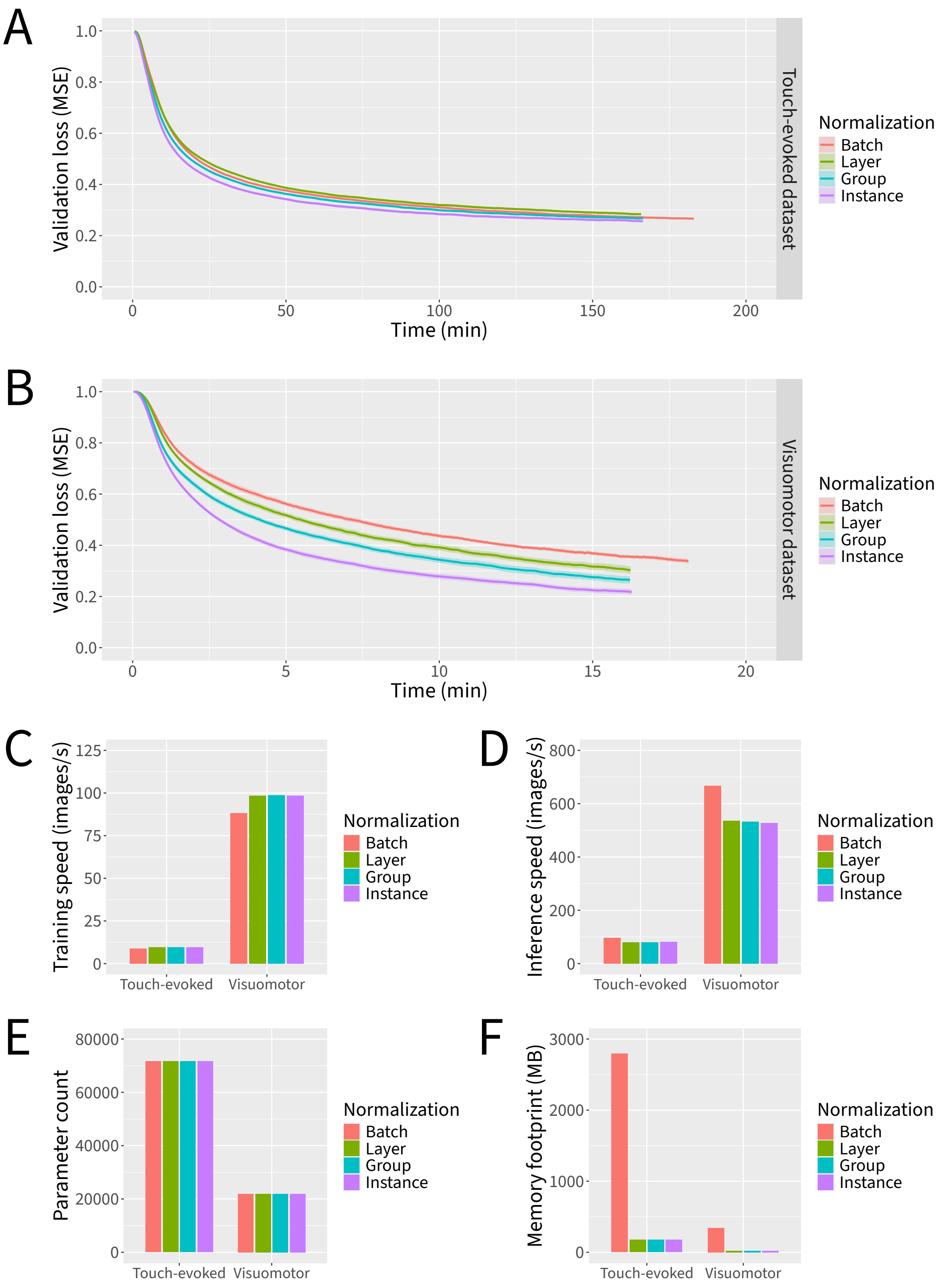


**Supplementary Figure 2. Ablation experiment for the "Swap" micro-architectural design step.** Neural networks were trained for 6000 parameter updates for each dataset and type of normalization. (**A**) Loss curves for the touch-evoked response dataset. (**B**) Loss curves for the visuomotor response dataset. (**C**) Training speeds. (**D**) Inference speeds. (**E**) Parameter counts. (**F**) Theoretical minimum memory footprints. Data represent the mean plus and minus the standard error of the mean of 10 independent experiments in which datasets were partitioned and neural network weights were initialized using different random seeds.

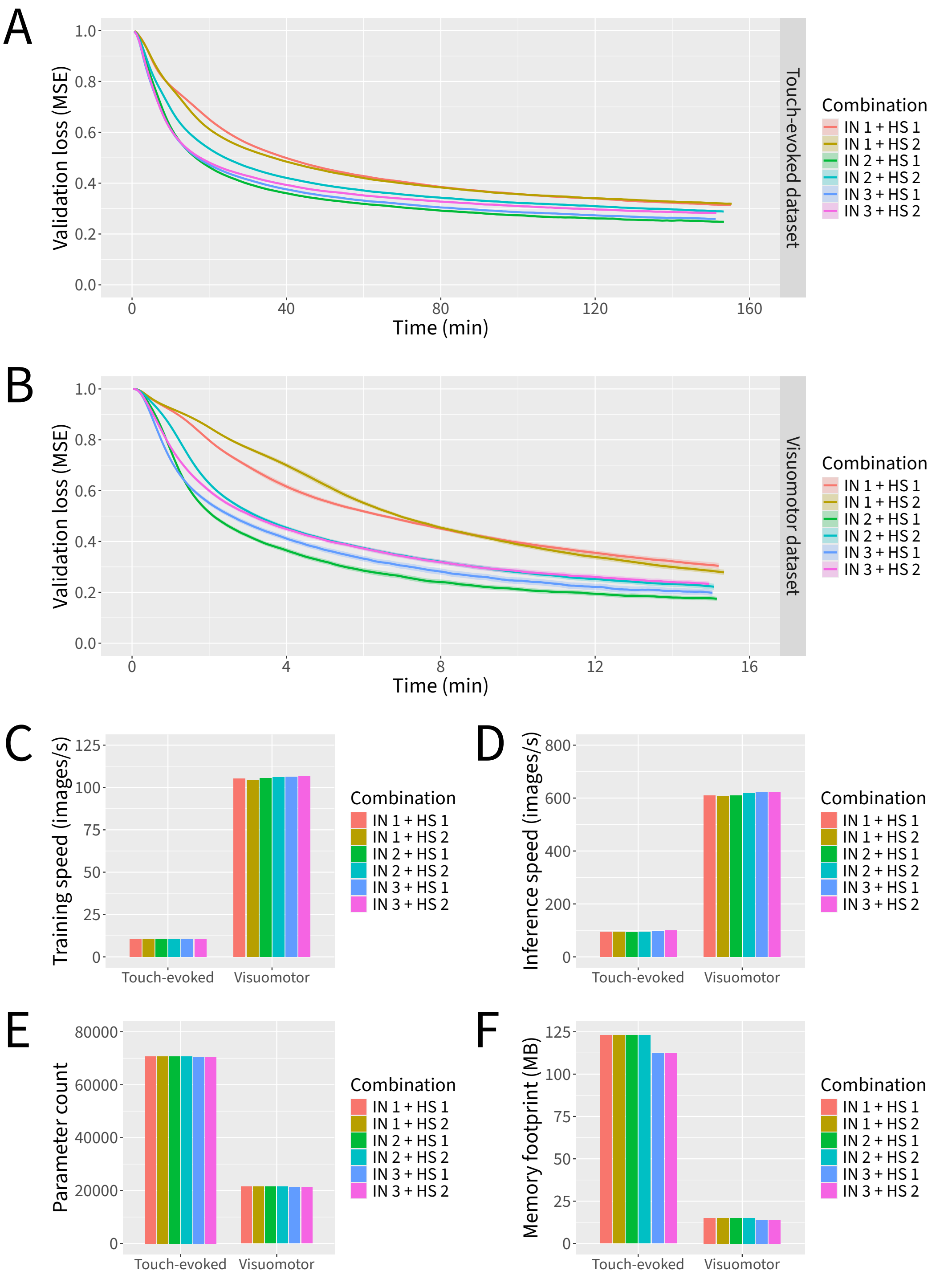


**Supplementary Figure 3. Ablation experiment for the "Drop" micro-architectural design step.** Neural networks were trained for 6000 parameter updates for each dataset and combination of Instance Normalization and Hard Swish layers. (**A**) Loss curves for the touch-evoked response dataset. (**B**) Loss curves for the visuomotor response dataset. (**C**) Training speeds. (**D**) Inference speeds. (**E**) Parameter counts. (**F**) Theoretical minimum memory footprints. Data represent the mean plus and minus the standard error of the mean of 10 independent experiments in which datasets were partitioned and neural network weights were initialized using different random seeds. IN: Instance Normalization; HS: Hard Swish.
